# Mapping the cellular and molecular organization of mouse cerebral aging by single-cell transcriptome imaging

**DOI:** 10.1101/2022.09.14.508048

**Authors:** William E. Allen, Timothy R. Blosser, Zuri A. Sullivan, Catherine Dulac, Xiaowei Zhuang

## Abstract

The cellular diversity and complex organization of the brain have hindered systematic characterization of age-related changes in its cellular and molecular architecture, limiting our ability to understand the mechanisms underlying its functional decline during aging. Here we generated a high-resolution cell atlas of brain aging within the frontal cortex and striatum using spatially resolved single-cell transcriptomics and quantified the changes in gene expression and spatial organization of major cell types in these brain regions over the lifespan of mice. We observed substantially more pronounced changes in the composition, gene expression and spatial organization of non-neuronal cells over neurons. Our data revealed molecular and spatial signatures of glial and immune cell activation during aging, particularly enriched in subcortical white matter, and identified both similarities and notable differences in cell activation patterns induced by aging and systemic inflammatory challenge. These results provide critical insights into age-related decline and inflammation in the brain.

## INTRODUCTION

The mammalian brain exhibits remarkable stability over periods ranging from years to decades (Yankner et al., 2008). Due to the brain’s limited regenerative abilities, neurons must faithfully perform their function for the entire lifetime of an animal. However, as the animals age, this longevity of neurons makes the brain sensitive to the accumulation of damage over time (Yankner et al., 2008). This neuronal damage, in combination with age-dependent changes in non-neuronal cells that support neural circuit function, is thought to cause the decline of brain function, increased sensitivity to damage, and drastic increase in the prevalence of neurodegenerative disorders (Bishop et al., 2010; Lindenberger, 2014; Yankner et al., 2008). Decades of research have provided rich insights into the molecular and cellular factors associated with brain aging, suggesting a complex process that so far escapes full understanding.

One prominent hypothesis suggests that changes in neuronal and synaptic functions associated with age and neurodegeneration are the result of disruptions to the brain’s homeostatic environment (Labzin et al., 2018; Mosher and Wyss-Coray, 2014; Richard M. Ransohoff, 2016). Neurons are supported by a host of non-neuronal cells, including glial cells such as astrocytes and oligodendrocytes, immune cells such as microglia, and various vascular cells, each maintaining different aspects of the tissue homeostasis (Meizlish et al., 2021) to ensure proper brain function. For example, oligodendrocytes myelinate axons and provide metabolic support to neurons; astrocytes provide trophic and ionic support to neurons and modulate synaptic function; and microglia provide immune surveillance, synaptic pruning, as well as debris removal by phagocytosis (Alves De Lima et al., 2020; Andreone et al., 2015; Croese et al., 2021; Ben Haim and Rowitch, 2016; Hammond et al., 2018; Li and Barres, 2018; Monje, 2018; Sofroniew, 2020). Brain injury, infection, and neurodegeneration have been shown to trigger inflammatory activation of these resident non-neuronal cell types and recruit peripheral immune cells, resulting in both protective and deleterious effects for neighboring neurons (Bohlen et al., 2019; Croese et al., 2021; Hammond et al., 2018; Labzin et al., 2018; Sofroniew, 2020).

Recent transcriptomic studies of normal brain aging (Almanzar et al., 2020; Benayoun et al., 2019; Schaum et al., 2020; Ximerakis et al., 2019) and neurodegenerative disease (Chen et al., 2020b; Grubman et al., 2019; Lau et al., 2020; Mathys et al., 2019), as well as studies focusing on specific non-neuronal cell types such as astrocytes (Boisvert et al., 2018; Clarke et al., 2018; Habib et al., 2020), microglia (Hammond et al., 2019; Olah et al., 2018), and endothelial cells (Chen et al., 2020a), have further highlighted a role for inflammatory activation and the disruption of non-neuronal cell states in aging-related decline. In particular, reactive states that are typically triggered in both microglia and astrocytes during infection or injury and that disrupt the normal homeostatic functions of these cell types, emerge naturally over the course of normal aging, even in the absence of overt neurodegenerative diseases.

While these studies suggest broad age-related disruptions to brain homeostasis that manifest in a variety of cell types, they also raise many questions. For example, how do the composition, molecular signatures, and spatial organization of different cell types and states in the brain change over aging and how do these changes relate to age-induced inflammatory activation? How are activated cells spatially distributed and does this activation depend on particular environmental factors and specific cell-cell communications? How does age-induced inflammation relate to systemic inflammation? Answering these questions is challenging as the brain’s enormous cellular and molecular complexity has so far prevented a comprehensive understanding of the changes in brain architecture over an animal’s lifetime.

Here, we performed a systematic characterization of the changes in molecular signatures and spatial organizations of cells during brain aging by using an experimental approach that combines single-nucleus RNA sequencing (snRNA-seq) (Habib et al., 2017) with a single-cell transcriptome imaging method, multiplexed error-robust fluorescent in situ hybridization (MERFISH) (Chen et al., 2015). This approach allowed us to profile gene expression and identify cell types in the mouse frontal cortex and stratum, thus generating a spatially resolved cell atlas of these regions at various ages during the lifespan of mice. This high-resolution cell atlas revealed age-related changes in both neurons and non-neuronal cells and uncovered molecular and spatial signatures of glial and immune cell activation during aging. Comparison of these changes with those induced by lipopolysaccharide (LPS) further revealed previously unknown differences in non-neuronal cell activation induced by aging and by systemic inflammatory challenge.

## RESULTS

### Spatially resolved single-cell transcriptomic profiling of the aging brain

We started with snRNA-seq measurements to probe the transcriptomic profiles of individual cells from the frontal cortex and striatum of mice at two different ages, 4-week and 90-week postnatal (**Figure 1A and 1B**). These brain regions have been previously shown to be susceptible to various age-related neurodegenerative diseases in humans (O’Callaghan et al., 2014; Seelaar et al., 2011). We sequenced ∼50,000 nuclei from these regions from two female animals at each age and performed unsupervised clustering analysis of the ∼80,000 cells that passed quality control (**Figure S1**).

**Figure 1:**
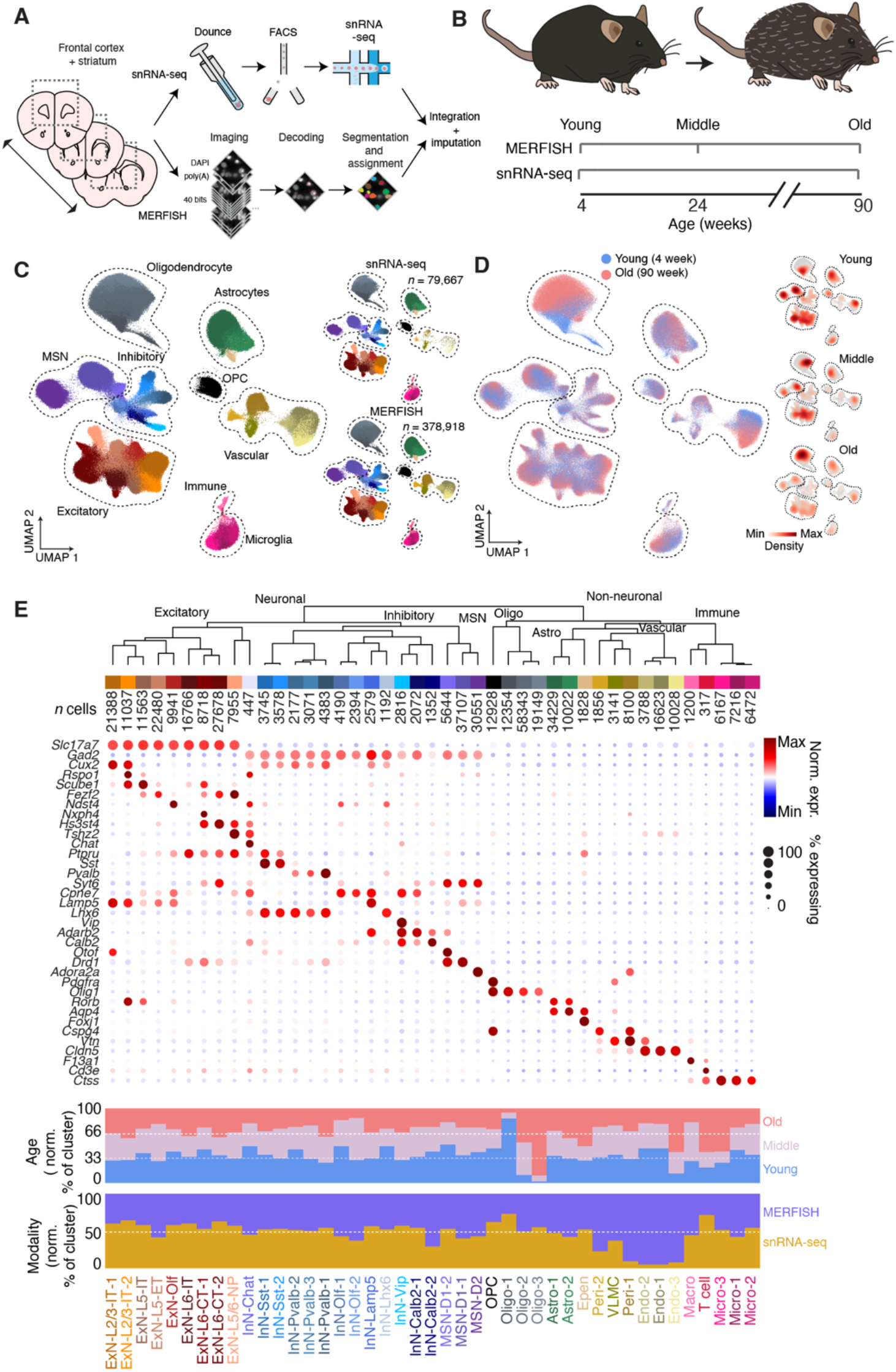
Spatially resolved single-cell transcriptomic profiling of the mouse frontal cortex and striatum over the animal’s lifespan. **(A)** Profiling of the mouse frontal cortex and striatum via integrated snRNA-seq and MERFISH analyses. **(B)** Sampling time points for snRNA-seq and MERFISH measurements across the lifespan of mice. **(C)** (Left) Uniform Manifold Approximation (UMAP) visualization of cells from all timepoints, from both snRNA-seq and MERFISH measurements. (Right) Separate UMAP visualization of cells measured by snRNA-seq (top) and MERFISH (bottom). Cells are colored by cell-type assignment. **(D)** (Left) As in (C) but with cells colored by age. Only two of the time points (Young and Old) are shown. (Right) Individual UMAP plots, shown as the density of cells at each time point (Young; Middle age; Old) overlaid on total cell population (grey) across all three ages. **(E)** Molecularly defined cell types determined from integrated snRNA-seq and MERFISH clustering analysis. (Top) Dendrogram of the hierarchical relationship among clusters and number of measured cells per cluster. (Middle) Expression of marker genes, showing major marker genes for different cell types. (Bottom) Fraction of cells per cluster by age and by modality, normalized to sampling depth such that equal representation in each condition will have the same fraction.

We then selected two sets of genes for spatially resolved single-cell transcriptomic measurements by MERFISH based on the snRNA-seq results (**Figure 1A**): i) cell type markers that were differentially expressed between cell clusters determined by snRNA-seq; ii) aging-related genes that were differentially expressed between the two ages in individual cell clusters. In addition, we selected previously known cell-type marker genes that define major neuronal, glial, and immune cell types and genes previously reported to be highly upregulated in various cell types during aging. Together, these resulted in a total of 212 cell-type markers and 204 aging-related genes, which we imaged in the same tissue sections through two back-to-back MERFISH runs, each with a 20-bit barcoding scheme (**STAR Methods** and **Table S1**).

We performed MERFISH imaging of these 416 genes in the frontal cortex and striatum across three different ages, 4-week (young), 24-week (middle-age), and 90-week (old) postnatal, including 3 – 5 female animals at each age, measuring a total of ∼400,000 cells after quality control (**Figure 1B**). The expression levels of individual genes measured by MERFISH showed good correlation with results from bulk RNA sequencing and were highly reproducible between biological replicates (**Figure S2A-C**). We then co-embedded the MERFISH and snRNA-seq data and performed an integrated clustering analysis across these two data modalities (**Figure 1C and 1D**), which showed good correspondence with the clustering results from the snRNA-seq alone (**Figure S2D**). The integrated analysis resulted in a total of 43 neuronal and non-neuronal cell types (**Figure 1E**). Compared to our previous MERFISH results in the mouse cortex (Zhang et al., 2021) and striatum (Chen et al., 2021), cell clusters were dissected here at a lower granularity to capture age-related changes of major cell types. The neuronal clusters included layer-specific excitatory neuronal cell types (ExN) in the cortex (ExN-L2/3-IT, ExN-L5-IT, ExN-L5-ET, ExN-L5/6-NP, ExN-L6-IT, and ExN-L6-CT), inhibitory neuronal cell types (InN) in the cortex marked by canonical inhibitory neuronal markers (*Sst*, *Pvalb*, *Lamp5*, and *Vip*), excitatory and inhibitory neurons in the subcortical olfactory areas (ExN-Olf and InN-Olf), and *Drd1+* (D1) or *Drd2+* (D2) medium spiny neurons (MSN) and *Lhx6+* or *Chat+* interneurons in the striatum, as well as spatially dispersed *Calb2+* interneurons. The non-neuronal clusters include oligodendrocytes (Oligo), oligodendrocyte precursor cells (OPC), astrocytes (Astro), ependymal cells (Epen), pericytes (Peri), vascular leptomeningeal cells (VLMC), endothelial cells (Endo), and microglia (Micro), macrophages (Macro), and T cells. snRNA-seq and MERFISH data co-embedded well with each other and the vast majority of cell clusters were well represented in both datasets with high correspondence in gene expression between the clusters from each dataset (**Figure 1C and 1E; Figure S2E**). However, we noted that some vascular cell types (pericytes and endothelial cells) were poorly sampled by snRNA-seq (**Figure 1E**).

In addition to leveraging both snRNA-seq and MERFISH data for cell type identification, this integration also allowed us to impute genome-wide expression profiles for individual cells measured by MERFISH using the transcriptomic profiles of neighboring snRNA-seq cells in the gene-expression space (**STAR Methods**). As a validation for the imputation results, the spatial distributions of the genes determined from the imputation results showed good agreement with both the results directly measured by MERFISH (for genes included in the MERFISH gene panel) and the results from Allen brain in situ hybridization atlas (for genes not included in the MERFISH gene panel) (**Figure S3**).

### Age-related changes in cell state and composition

We next analyzed how the cellular composition of these brain regions changed over aging based on the *in situ* MERFISH data. The neuronal clusters did not exhibit significant changes in abundance across the three ages (**Figure 2A**). By contrast, several non-neuronal cell types exhibited substantial age-dependent changes in the overall abundance of the cell type and/or the relative proportions of cells among different subtypes or states within the cell type (**Figure 1E; Figure 2A and 2B**). In particular, the abundance of oligodendrocytes increased and that of the OPCs decreased substantially as the animal aged (**Figure 2A**). Of the three subtypes or states of oligodendrocytes, Oligo-1 was predominant in young animals and diminished to nearly non-existent in middle-aged and old animals, Oligo-2 was predominant in middle-aged animals and decreased in abundance in old animals, while a third cluster Oligo-3 emerged in old animals (**Figure 2B**). The aging-related cluster Oligo-3 exhibited substantially upregulated expression of *C4b* (**Figure 2C**), a complement protein of the innate immunity system, and interleukin 33 (*Il33*) (**Figure 2C**), a cytokine involved in inflammatory and innate immune response (Molofsky et al., 2015) and previously shown to be upregulated in oligodendrocytes in the aging brain (Ximerakis et al., 2019). These results suggest an initial maturation and proliferation of oligodendrocytes, likely a result of late-stage development, followed by inflammatory activation of matured oligodendrocytes with aging. Microglia, endothelial cells, and astrocytes did not exhibit substantial overall abundance change but showed age-dependent shift in population among subtypes or states. For example, Micro-1 and Endo-1 were enriched in young animals; Micro-3 and Endo-3 were enriched old animals; Astro-2 showed increased abundance in old animals (**Figure 2B**). These aging-related cell subtypes or states exhibit upregulation of genes (e.g., *B2m* and *Trem2* in Micro-3, *Xdh* in Endo-3, and *Gfap* and *C4b* in Astro-2) (**Figure 2C**), some of which have been previously shown to be enriched in microglia, endothelial cells, and astrocytes activated by inflammation and/or aging (Boisvert et al., 2018; Chen et al., 2020a; Clarke et al., 2018; Hammond et al., 2019; Liddelow et al., 2017; Ximerakis et al., 2019). Consistent with previously observed T cell infiltration into the aging brain (Dulken et al., 2019), we also observed a substantial increase in the abundance of T cells in old animals (**Figure 2A**), although the change did not reach statistical significance due to the small total number of cells detected for this rare cell type.

**Figure 2:**
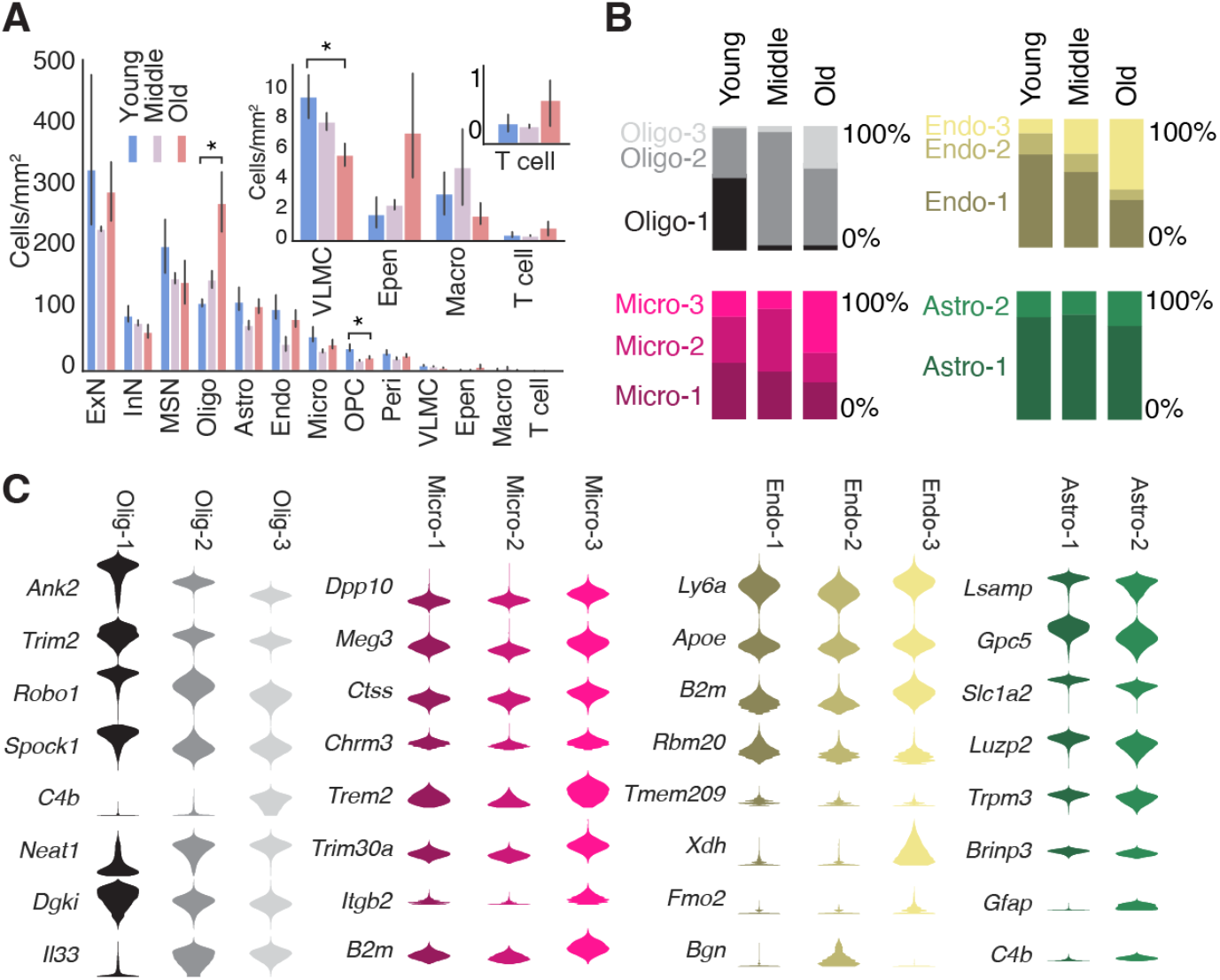
Changes in cell-type composition of the mouse frontal cortex and striatum across ages. **(A)** Density of different major cell types (in cells/mm^2^) across the three ages (young: 4 weeks; middle: 24 weeks; old: 90 weeks). Inset shows magnified view of lower abundance cell types. * indicates FDR-adjusted *P*-value < 0.05 for independent sample *t*-test in difference in density between young and old animals. Error bars: 95% confidence interval. **(B)** Fraction of cells that belong to different subtypes or states of oligodendrocytes, microglia, endothelial cells, and astrocytes across different ages. **(C)** Violin plot of expression of example genes that change expression across different subtypes or states of oligodendrocytes, microglia, endothelial cells, and astrocytes.

### Age-related changes in the spatial organization of individual cell types

The *in situ* cell-type identification by MERFISH further allowed us to map the spatial organization of individual cell types across different ages. To visualize the overall spatial organization of cells, we performed hierarchical clustering of cells based on the cell composition in their spatial neighborhood (**STAR Methods**) and the resulting spatial clusters naturally segmented the imaged brain regions into several subregions that corresponded to known anatomical features, including the pia, different cortical layers, corpus callosum, striatum, ventricle, and subcortical olfactory regions (**Figure 3A**). As expected, different excitatory clusters adopted laminar distributions in the cortex and medium spiny neurons were localized to the striatum (**Figure 3A and 3B**); oligodendrocytes were enriched in the corpus callosum, vascular leptomeningeal cells and a specific pericyte cluster (Peri-2) were enriched in the pia, and ependymal cells were localized around the ventricle, whereas OPCs, astrocytes, microglia and endothelial cells were distributed largely uniformly throughout the imaged regions (**Figure 3A and 3B**).

**Figure 3:**
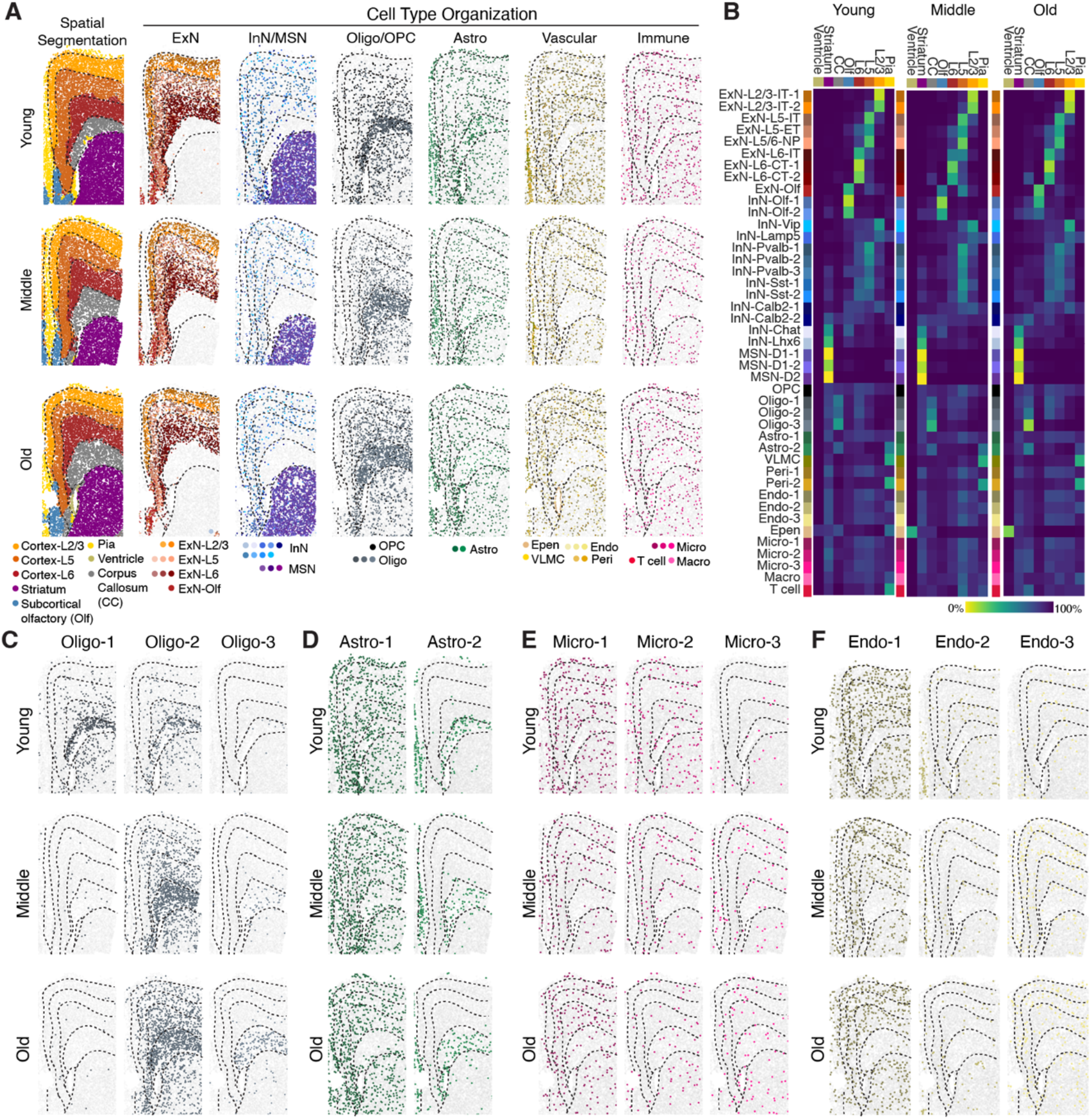
Spatial organization of cells in the mouse frontal cortex and striatum across ages. **(A)** (Left) Spatial segmentation of anatomical regions. (Right) Spatial organization of major cell types at the three different ages, colored by cluster identity. Dashed lines outlining anatomical regions were manually traced from spatial segmentation for visualization purpose. **(B)** Fraction cells resided in individual anatomical regions for each cell cluster at the three different ages. CC: corpus callosum. Olf: Subcortical olfactory areas. The lower abundance of ependymal cells in younger animals may be due to classification as molecularly similar astrocytes or loss of ventricle surface during tissue sectioning. **(C–F)** Spatial organizations of oligodendrocyte (C), astrocyte (D), microglial (E), and endothelial clusters at different ages.

Interestingly, although the overall spatial organization of neuronal cell types appeared similar across different ages (**Figure 3A and 3B**), some non-neuronal cell clusters showed changes in anatomical enrichment with age. For example, the oligodendrocyte cluster that emerged in old animals (Oligo-3) was located nearly exclusively in the corpus callosum, whereas cells from the Oligo-1 and Oligo-2 clusters, albeit being enriched in the corpus callosum, could be found throughout the imaged regions (**Figure 3C**). Likewise, astrocytes belonging to the aging-related Astro-2 cluster were localized to the corpus callosum, whereas cells from Astro-1 adopted a complementary distribution that became depleted in corpus callosum in adult and aged animals (**Figure 3D**). Not all cell types that exhibited age-dependent shift in subtypes or states showed spatial heterogeneity – different microglial and endothelial clusters were more-or-less evenly distributed throughout all anatomical regions (**Figure 3E and 3F**).

In addition, we observed that certain non-neuronal cell types exhibited a tendency to be spatially colocalized, which further increased with age. Specifically, vascular cells (endothelial cells, pericytes and vascular leptomeningeal cells) showed a significant tendency to be proximal to each other and this tendency increased with age (**Figure S4**). Moreover, macrophages tended to be enriched near vascular cells and this tendency also increased with age (**Figure S4**). A similar trend was also observed for microglia, albeit to a lesser degree (**Figure S4**).

### Age-related changes in the gene expression profiles of individual cell types

Overall, the above results showed that both the composition and the spatial organization of cells exhibited aging-induced changes primarily in non-neuronal cell types. Next, we examined how gene expression profiles of individual cell types changed with age. To this end, we determined the number of genes that were differentially expressed across different ages in individual neuronal and non-neuronal cell types based on the gene expression profiles determined by snRNA-seq (**Figure 4A**). While essentially all cell types had at least some genes that were differentially expressed over aging, non-neuronal cell types tended to exhibit a greater number of age-dependent differentially expressed genes (**Figure 4A**). The majority of the age-dependent genes were differentially expressed in a cell-type-specific manner, with relatively few genes broadly differently expressed across all cell types (**Figure 4B**). Gene Ontology (GO) and KEGG (Kyoto Encyclopedia of Genes and Genomes) enrichment analysis showed that genes upregulated with age in neurons, in particular in inhibitory neurons, were enriched in pathways associated with neurodegenerative diseases, oxidative response and mitochondria function, whereas genes upregulated with age in non-neuronal cell types tended to be associated with inflammatory and immune response (**Figure 4C**), consistent with the previous observations of broad upregulation of oxidative stress and immune pathways in the aging brain (Lu et al., 2004; Almanzar et al., 2020; Benayoun et al., 2019; Ximerakis et al., 2019). Specifically, these age-upregulated genes in non-neuronal cells included cytokines (e.g. *Il33* and *Il18* in oligodendrocytes), complement proteins (e.g. *C4b* in astrocytes and oligodendrocytes), and proteins involved in interferon response (e.g. *Ifit3* and *Ifitm3* in ependymal cells and pericytes) (**Figure 4B**).

**Figure 4:**
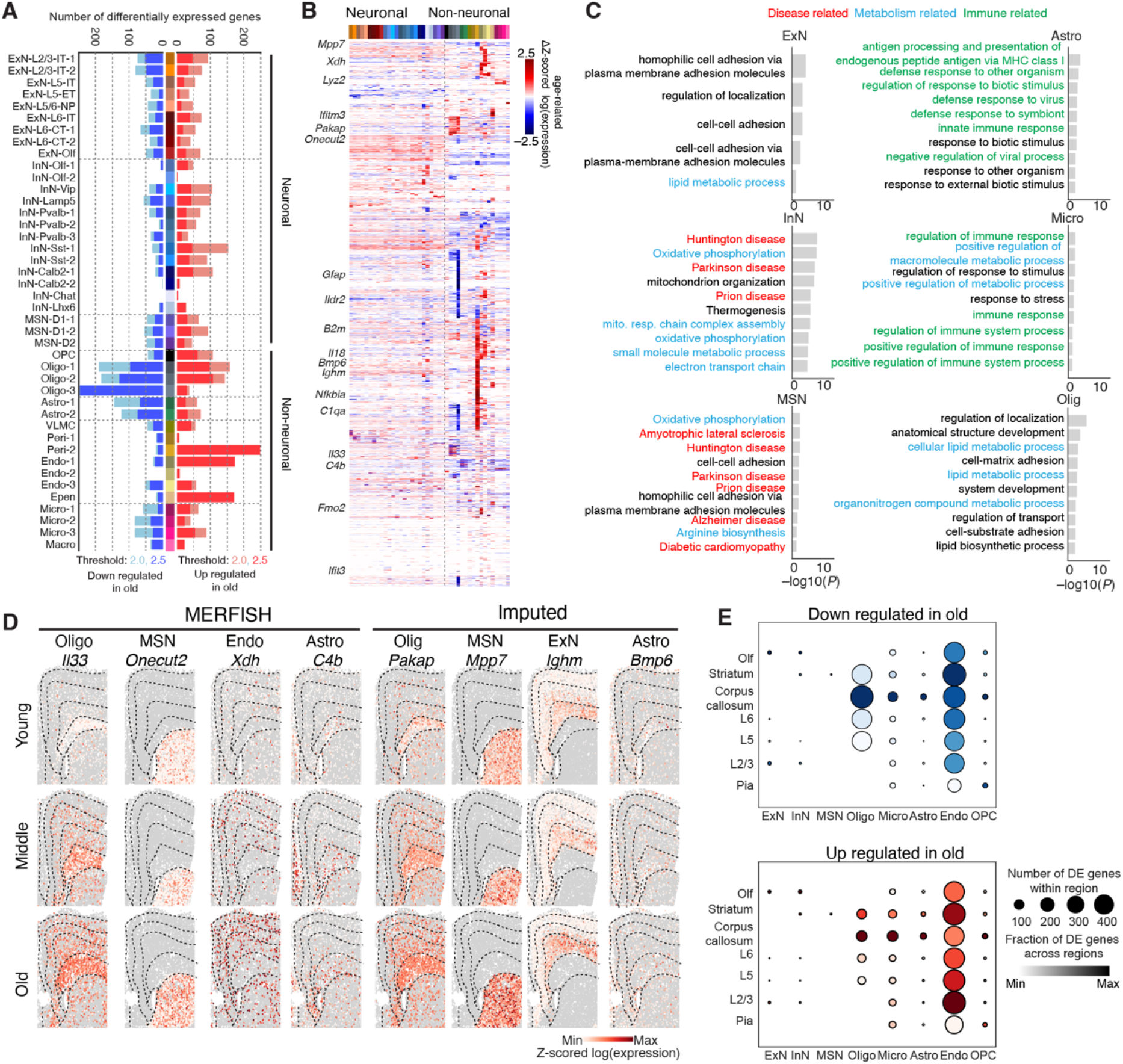
Cell-type-specific and spatially localized molecular signatures of brain aging. **(A)** Number of genes differentially expressed between young and old animals in individual cell clusters, with genes upregulated with age shown in red bars and downregulated with age shown in blue bars. Differentially expressed (DE) genes were defined as genes with age-related change in log(gene expression) > 2 (light colored bars) or > 2.5 (dark colored bars) and FDR-adjusted *P*-value < 0.05 between the two ages. **(B)** Age-related change in Z-scored log(gene expression) between young and old animals for DE genes in different cell clusters. **(C)** –log_10_(*P*-value) of enrichment for Gene Ontology (GO) Biological Process terms and Kyoto Encyclopedia of Genes and Genomes (KEGG) terms enriched among DE genes with an age-related change in log(gene expression) > 2 and FDR-adjusted *P*-value < 0.05 between the two ages. Only GO or KEGG terms with *P*-value < 0.05 are listed and when the number of terms exceeds 10, only top 10 terms are listed for each major cell class. **(D)** Spatial maps of examples of DE genes across the three different ages. **(E)** Quantification of the number of DE genes for each major cell type as a function of spatial location, using imputed gene expression data. DE genes with an age-related change in log(gene expression) > 2 and FDR-adjusted *P*-values < 0.05 are considered. Size of dot indicates total number of DE genes for a particular cell type within a region, color shade of dot indicates the fractional number of DE genes relative to the maximum value across all regions.

Imputation of the genome-wide expression profiles of the cells measured by MERFISH allowed us the determine the spatial distributions of all genes across different ages. Many of the age-upregulated genes exhibited specific spatial patterns, such as being highly enriched in the corpus callosum, specific cortical layers, striatum or other anatomical regions (see examples in **Figure 4D**). Using these imputed genome-wide expression profiles, we systematically quantified the number of genes differentially expressed over aging for each major cell type in individual anatomical regions. We observed substantial spatial heterogeneity in the total number of genes upregulated or downregulated with age even among cells of the same type (**Figure 4E**). In particular, oligodendrocytes, astrocytes, and microglia all exhibited the greatest number of differentially expressed genes with age in the white matter of the corpus callosum, relative to other anatomical regions. Endothelial cells, on the other hand, had more age-upregulated or - downregulated genes in the striatum and specific cortical layers, with more genes upregulated in upper layers.

To further investigate age-related changes in gene expression in non-neuronal cell types, we performed gene-gene correlation analysis to identify groups of genes whose expression showed correlated variations with each other and hence likely belong to the same gene regulatory networks. Within each cell type, we determined pair-wise Pearson correlation coefficient of gene expression across all cells measured at all ages for any pair of genes. Such gene-gene correlation matrices for oligodendrocytes, astrocytes, and microglia revealed, for each cell type, many groups of genes that showed correlated expression, which we referred to as gene modules (**Figure S5**). Many of these gene modules showed up- or down-regulation in expression with age (**Figure S5**). GO or KEGG term analysis showed that many of these modules were related to development or immune response, often capturing cell-type specific functions. For example, microglia module 20 and oligodendrocyte module 23 were upregulated with age and enriched for terms related to inflammatory and immune response, such as “innate immune response”, “response to cytokine”, and “cellular response to interferon beta”, whereas oligodendrocyte module 12 was enriched for terms related to myelination, such as “axon ensheathment” and “nervous system development” (**Figure S5**). These results suggest the presence of specific gene regulatory networks that function in a cell-type-specific and age-dependent manner, whose functional annotation suggests specific biological functions of those cell types (e.g. myelination) or responses to external stimuli (e.g. cytokines).

### Age-dependent activation of glial and immune cells

For microglia and astrocytes, we observed that the genes highly upregulated with age overlapped substantially with genes that have been previously reported as being upregulated in the activated (or ‘reactive’) state of these cell types (e.g. *Gfap* and *C4b* for astrocytes; *B2m* and *Lyz2* for microglia) (Clarke et al., 2018; Hammond et al., 2019; Keren-Shaul et al., 2017; Liddelow et al., 2017). These activated astrocytes and microglia have been observed in both healthy and diseased brains, often responding to brain injury, inflammation, or degeneration, and have been shown to have protective or deleterious effects on neighboring neurons depending on the specific type and level of activation (Bohlen et al., 2019; Croese et al., 2021; Hammond et al., 2018; Labzin et al., 2018; Mhatre et al., 2015; Sofroniew, 2020). Microglial and astrocytic activation has been reported in aged rodent and human brains (Boisvert et al., 2018; Clarke et al., 2018; Habib et al., 2020; Hammond et al., 2019; Olah et al., 2018), but how such activation depends on the spatial context remains unclear.

To quantify the activation of these cell types and determine the spatial distributions of activated cells, we scored the activation levels of astrocytes and microglia imaged by MERFISH using genes previously shown to be specific for activated cells (**STAR Methods**). The activation scores for both astrocytes and microglia increased on average with age (**Figure 5A**). The specific subtypes or states of astrocytes and microglia enriched in old animals (Astro-2 and Micro-3) had higher activation scores than the other subtypes or states (**Figure 5B**).

**Figure 5:**
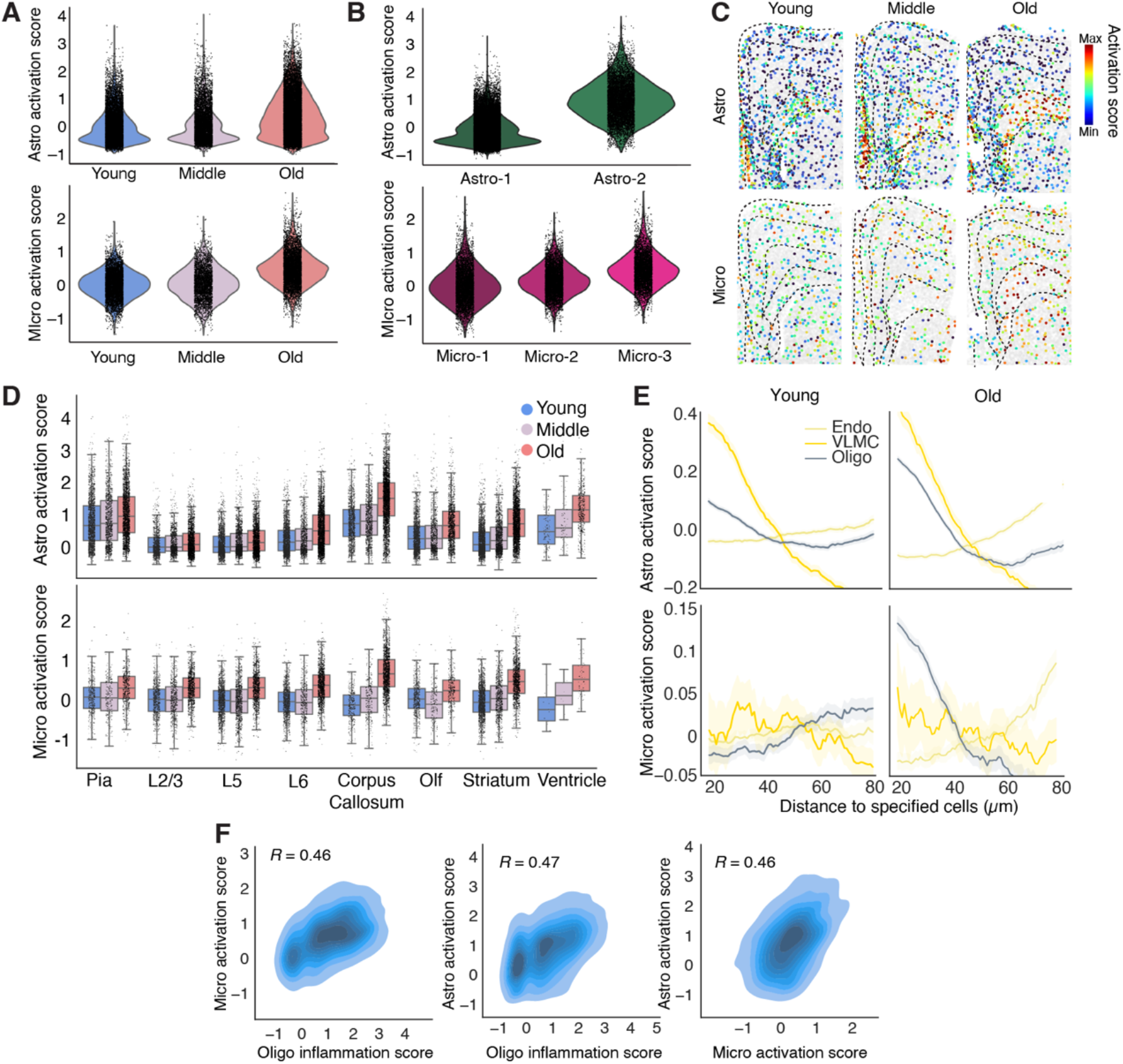
Spatially heterogeneous and cell-type-specific inflammatory activation signatures of brain aging. **(A)** Activation scores of all astrocytes and microglia across the three different ages. Activation score is defined as the summed expression of a cell-type-specific subset of gene related to inflammatory activation, relative to background of randomly selected genes (**STAR Methods**). **(B)** Activation scores of specific astrocyte and microglia clusters. **(C)** Spatial maps of activation scores of astrocytes and microglia across the three different ages. Cell are colored by activation scores. **(D)** Quantifications of per-cell activation scores of astrocytes and microglia in different anatomical regions across three ages. **(E)** Average activation scores of astrocytes and microglia as a function of distance from neighboring oligodendrocytes, vascular leptomeningeal cells (VLMCs), and endothelial cells. **(F)** (Left) Correlation plots of activation scores of each microglial cell in corpus callosum versus the average inflammation scores of oligodendrocytes within 30 µm of that microglial cell. (Middle) Same as (Left) but for astrocytes and oligodendrocytes. (Right) Same as (Left) but for astrocytes and microglia. Pearson correlation coefficients *R* are given in the plots.

Notably, astrocyte and microglia exhibited distinct spatial signatures in their activation patterns. Astrocytes showed pronounced spatial heterogeneity in activation, with the highest level of activation in the corpus callosum, as well as relatively strong activation in the striatum and near the ventricle and pia, but minimal activation in the cortex (**Figure 5C and 5D**). This spatial pattern was already apparent in young animals and became more pronounced with aging. Microglia activation, on the other hand, was more uniform across different regions, with hardly any spatial heterogeneity in young animals (**Figure 5C and 5D**). As the animal aged, microglia activation level increased more-or-less uniformly across different regions, except that the corpus callosum showed a notably higher level of activation (**Figure 5C and 5D**).

Because microglia and astrocytes may be directly or indirectly activated by pro-inflammatory cytokines and chemokines that circulate in the blood (Pluvinage and Wyss-Coray, 2020) or are released by brain-infiltrating immune cells (Croese et al., 2021), and we observed enrichment of macrophages near vascular cells in old animals (**Figure S4**), we further examined whether the activation levels of astrocytes and microglia depended on their distance to vascular cells that separate the bloodstream (e.g. endothelial cells) or cerebrospinal fluid (e.g. vascular leptomeningeal cells (VLMCs)) from the interior of the brain. In addition, since we observed that several genes involved in inflammatory response and innate immune signaling (*Il18*, *Il33*, and *C4b*) were upregulated in oligodendrocytes (**Figure 4B**), we also examined the dependence of astrocyte and microglia activation on the distance to oligodendrocytes.

Again, these dependencies were notably different between astrocytes and microglia. Astrocyte activation exhibited a strong dependence on the proximity to VLMCs in both young and old animals, and a dependence on the proximity to oligodendrocytes that increased substantially with age (**Figure 5E**). Only the other hand, microglia did not show any preferential activation near vascular cells, but aging-induced activation of microglia showed a strong dependence on their proximity to oligodendrocytes (**Figure 5E**). Moreover, we scored the inflammation level of oligodendrocytes using the expression levels of *Il33*, *Il18*, and *C4b*, and observed that within the corpus callosum, the aging-induced activation levels of astrocytes and microglia were correlated with the inflammation level of nearby oligodendrocytes (**Figure 5F**), suggesting that the dependence on the proximity to oligodendrocytes was not simply a reflection of stronger activation of astrocytes and microglia in the corpus callosum but are likely related to the inflammatory response of oligodendrocytes over aging. The activation levels of astrocytes and microglia in the corpus callosum were also correlated with each other (**Figure 5F**).

The above results suggest multiple different mechanisms of non-neuronal cell activation, two of which showed strong spatial dependence: 1) activation of astrocytes near the surface of the vascular structures separating the cerebrospinal fluid and the brain, potentially caused by factors derived from cerebrospinal fluid; 2) activation of microglia and astrocytes near oligodendrocytes in corpus callosum, potentially caused by pro-inflammatory factors expressed by oligodendrocytes. Supporting this notion, the molecular signatures of activated astrocytes near the pia were different from those in the corpus callosum (**Figure S6**). Interestingly, only the second mechanism was aging specific.

### Activation of glial and immune cells in response to systemic inflammatory challenge

The activation of astrocytes and microglia with age, reminiscent of brain inflammation, raises an interesting question as to how these age-related states compare with those induced by systemic inflammation. Peripheral administration of LPS is widely used to model brain inflammation associated with neurodegenerative disease (Ribeiro et al., 2019). Although LPS itself is thought not to cross the blood-brain barrier, systemic release of cytokines and chemokines by peripheral immune cells upon LPS administration can broadly activate microglia and astrocytes throughout the brain (Clarke et al., 2018; Qin et al., 2007), as well as induce fever (Elmquist et al., 1996).

We injected mice at the three ages (4-week, 24-week, and 90-week postnatal) with LPS (**Figure 6A**), sacrificed the animals 24 hours after LPS injection, and performed MERFISH measurements using our cell type and aging gene panels. ∼350,000 cells passed quality control analysis, and we classified these cells by integrating the LPS dataset with the normal brain MERFISH dataset described earlier and transferred cell-type annotations without re-clustering (**Figure 6B; Figure S7A**).

**Figure 6:**
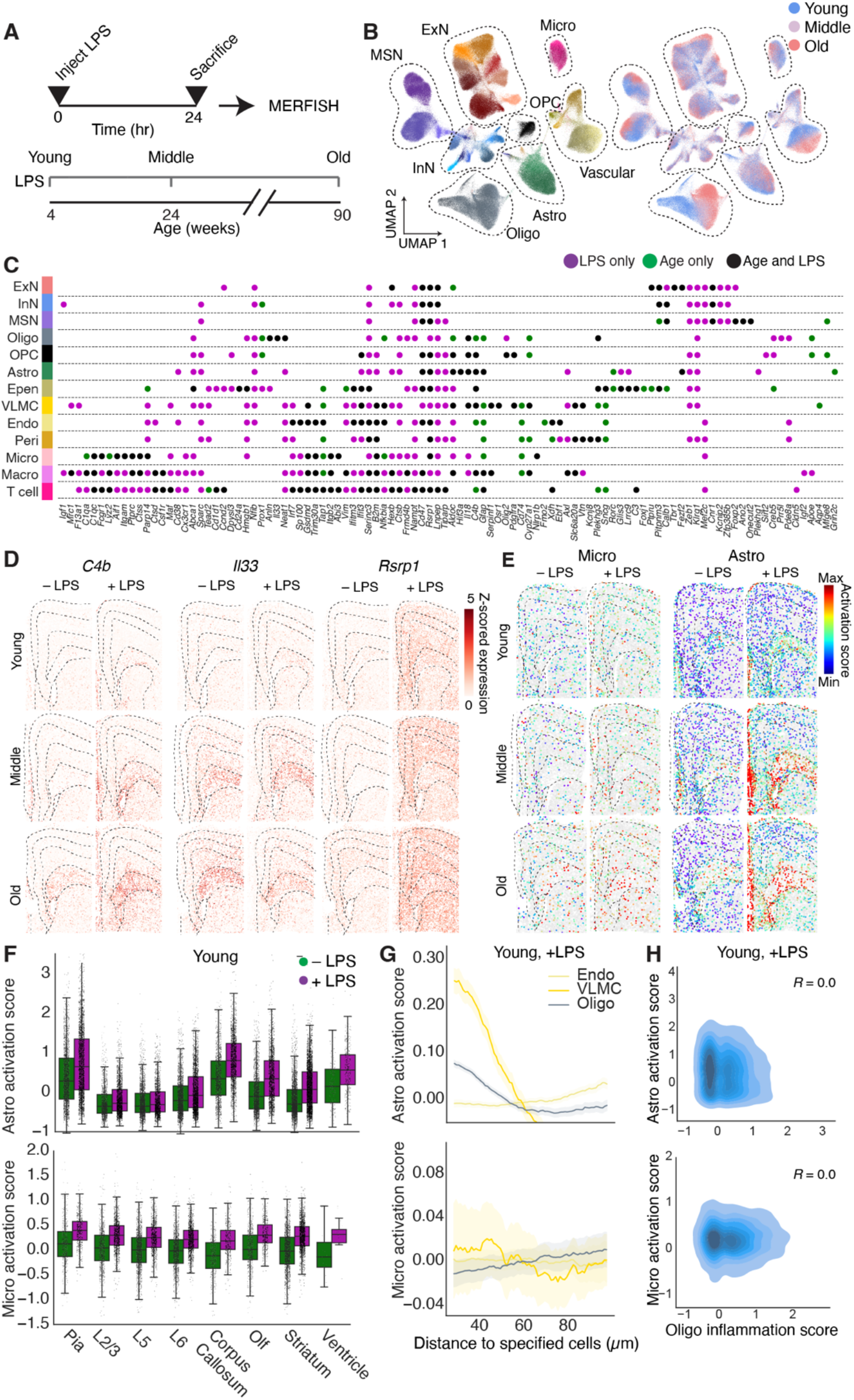
Gene expression changes and activations of cells in response to systemic inflammatory challenge. **(A)** Experimental paradigm for systemic inflammatory challenge with lipopolysaccharide (LPS). **(B)** UMAP visualization of cells colored by cell types (left) or age (right) measured by MERFISH. **(C)** Change in gene expression in response to LPS only (purple), aging only (green), or both LPS and aging (black) for different cell types measured by MERFISH. Only genes with age- or LPS-related change in Z-scored log(gene expression) > 2 in at least one condition for at least one cell type are shown. **(D)** Spatial maps of example genes that are upregulated over aging and upon LPS treatment. **(E)** Spatial maps of activated microglia and astrocytes across the three different ages, with and without LPS treatment. Cells are colored by activation scores. **(F)** Quantification of per-cell activation scores for microglia and astrocytes in different anatomical regions, in young mice with LPS treatment. **(G)** Activation scores of astrocytes and microglia as a function of distance from neighboring oligodendrocytes, VMLCs, and endothelial cells in young mice after LPS treatment. **(H)** Correlation of activation score of microglia and astrocytes and inflammation score of oligodendrocytes, as defined in **Figure 5F**, in young animals treated with LPS.

We observed a high degree of similarity between untreated and LPS-treated mice in terms of both the composition (**Figure S7A-C**) and the global spatial organization (**Figure S8A and S8B**) of the cell types. However, compared to untreated animals, young animals treated with LPS showed a substantially higher degree of enrichment of macrophages near vascular cells (**Figure S8C**), similar to that observed over the course of normal aging (**Figure S4**).

Notably, LPS induced substantial changes in the gene expression in a cell-type-specific manner and some of the upregulated genes overlapped with those observed over normal aging. To quantify these effects, we fit a regression model for each gene on young untreated and LPS-treated animals, or on untreated young and old animals, and compared the extent to which specific genes were upregulated across the two conditions (**Figure 6C; Figure S7D**). Here, we limited our analyses to the genes in the MERFISH panel and did not impute genome-wide expression because we did not perform scRNA-seq measurements on the LPS-treated animals. Nonetheless, these analyses provided interesting similarities and differences between aging- and LPS-induced changes. Many of the genes involved in innate immune response that were observed to be upregulated with age were also upregulated in responses to LPS (**Figure 6C; Figure S7D**). There was, however, substantial quantitative variations in the relative extent of upregulation under the two conditions. For example, *C4b* was highly upregulated over aging and further upregulated by LPS treatment, consistent with previous observations in astrocytes using bulk RNA-seq (Clarke et al., 2018); *Il33* was strongly upregulated with age whereas LPS treatment induced only very small additional upregulation of this gene; *Rsrp1* was more strongly upregulated in response to LPS and only weakly upregulate with age (**Figure 6D**). There was also a subset of immune-response related genes that were only upregulated under one of the two conditions, for example *Nfib, Sparc, Mef2c,* and *Zeb1* in response to LPS, and *Cd74* and *Mfge8* over aging (**Figure 6C**).

Because the MERFISH gene panel includes genes that are upregulated in reactive astrocytes and microglia in response to inflammation, the MERFISH data also allowed us to analyze the activation patterns of these cells under LPS treatment and compare with those induced by aging. LPS increased the activation of astrocytes and microglia in young and old animals (**Figure 6E**). Astrocytes were preferentially activated by LPS in or near the pia, corpus callosum, striatum, and ventricle but not the cortex, whereas microglia were largely uniformly activated by LPS across all regions (**Figure 6E and 6F**). Moreover, the activation of microglia by LPS did not depend on the proximity to oligodendrocytes or VLMCs, whereas the activation of astrocytes showed a strong dependence on the proximity to VLMCs and only a weak dependence to oligodendrocytes (**Figure 6G and 6H**). These results suggest interesting commonality and differences between age- and LPS-induced activations of non-neuronal cells: while both conditions induced spatially heterogeneous activation of astrocytes with particular enrichment near the cerebrospinal-fluid–brain barriers and dispersed activation of microglia, aging uniquely induced microglia activation, and potentially related increase in astrocyte activation, near oligodendrocytes in corpus callosum.

## DISCUSSION

How the brain ages and why these changes lead to functional decline are questions with major fundamental and practical significance, as one expects an increase in the prevalence of neurodegenerative diseases over the coming decades due to the aging global population. Many hypotheses have been proposed for the causes of brain function decline with age, ranging from changes in synaptic connectivity or physiology (Bishop et al., 2010), to senescence of glial and immune cells and the role of circulating inflammatory factors (Wyss-Coray, 2016). Previous transcriptomic studies have revealed widespread changes in cellular state with age, highlighting specific cell types and biological processes that might be involved in mediating age-related decline (Almanzar et al., 2020; Benayoun et al., 2019; Boisvert et al., 2018; Chen et al., 2020a; Clarke et al., 2018; Habib et al., 2020; Hammond et al., 2019; Olah et al., 2018; Schaum et al., 2020; Ximerakis et al., 2019). In particular, some of these studies have highlighted a role for increased inflammation as a key aspect of brain aging (Benayoun et al., 2019; Clarke et al., 2018; Hammond et al., 2019; Wyss-Coray, 2016). However, to understand how these changes may impact specific brain functions and gain insights into the mechanisms underlying age-related functional decline, it is crucial to characterize both the molecular and cellular signatures and the spatial locations of these changes within the brain.

To fill this gap, we used spatially resolved single-cell transcriptomic analysis to systemically uncover changes in the cellular composition, molecular signatures, and spatial organizations of brain cells in the mouse frontal cortex and striatum over the animal’s lifespan. By integrating snRNA-seq and MERFISH measurements, we generated a spatially resolved cell atlas of the aging brain with a genome-wide expression profile associated with each cell. Notably, we observed substantially more pronounced, and qualitatively different, age-induced changes in non-neuronal cells compared to neurons, and these changes in non-neuronal cell exhibited specific spatial patterns. This atlas provides a rich resource for understanding the changes in cell state associated with aging.

At the molecular level, many of the genes upregulated in non-neuronal cells during aging were related to activation of inflammatory pathways associated with innate immunity, while neuronal cell populations displayed different transcriptional changes, many of which related to neurodegenerative diseases, oxidative stress, and mitochondria functions. Immune cells and secreted factors such as cytokines are widely involved in the maintenance of tissue homeostasis across many organs (Meizlish et al., 2021). Hence, the observed upregulation of genes related to inflammation and innate immunity by immune and glial cells within the brain could be an indication of dysregulated tissue homeostasis that may broadly affect nervous system function. Furthermore, in each individual non-neuronal cell types, such as oligodendrocytes, astrocytes, and microglia, we observed dozens of gene modules that contained genes with correlated expression variation across cells, and many of these gene modules showed up- or down-regulated expression with aging. These modules suggest the presence of multiple, potentially interconnected, gene-regulatory networks related to aging.

While the cell composition and spatial organization of neurons were stable with age, we observed notable changes in the composition and spatial distributions of non-neuronal cells, with specific oligodendrocyte and astrocyte subtypes or states emerging in the corpus callosum of the aging brain. Interestingly, inflammatory activation of microglia and astrocytes during aging showed distinct spatial patterns: both cell types exhibited the strongest activation in the corpus callosum, a location that also showed strong inflammatory changes of oligodendrocytes, whereas astrocytes but not microglia showed increased activation near the pial surface. Overall, astrocyte activation appeared to be substantially more spatially localized than microglia activation. Taken together, these results highlight the white matter of the corpus callosum as a hotspot of age-associated inflammatory changes in the brain.

Previous MRI studies in humans have revealed that prefrontal white matter is highly susceptible to age-related reduction in volume (Gunning-Dixon et al., 2009), and that the degree of white matter changes are associated with cognitive decline (Gunning-Dixon and Raz, 2000). Electron microscopy studies of non-human primate brain aging have revealed major alterations specifically in the white matter, particularly in the disruption of myelin sheath structure (Peters, 2002). White matter microglia reactivity has also been related to aging (Safaiyan et al., 2021) and diseases, including, recently, SARS-CoV-2 induced long-term neurological impairment (Fernández-Castañeda et al., 2022). Expanding upon these pervious findings, our results suggest that changes in the oligodendrocytes and myelinated axons and their associated microglial and astrocytic reactivity in the white matter may be an important factor in age-associated cognitive deficits. Our observations that the activation levels of microglia and astrocytes in the corpus callosum are correlated with each other and both correlated with the inflammation level of oligodendrocytes marked by cytokines like Il33 further suggest potential molecular mechanisms underlying this inflammatory activation. In one scenario, the elevated expression of pro-inflammatory cytokine Il33 in aged oligodendrocytes may activate microglia through the Il33 receptors, which is known to be expressed in microglia (Vainchtein et al., 2018). In a second and potentially related model, myelin itself can activate microglia *in vitro* (Williams et al., 1994), and excessive myelin degradation can induce activation of phagocytosing microglia that become overloaded with myelin fragments (Safaiyan et al., 2016), in a process that depends on TREM2 signaling (Safaiyan et al., 2021). In both scenarios, activated microglia can in turn activate astrocytes through secretion of pro-inflammatory cytokines and complement proteins (Liddelow et al., 2017). Activated astrocytes and microglia may in turn exacerbate oligodendrocyte and myelin degeneration (Gibson et al., 2019; Liddelow et al., 2017).

Our result further showed that microglia and astrocyte activation induced by aging exhibited similarities but also significant differences to that induced by systemic inflammatory challenge. On the one hand, many of the same genes were upregulated by acute LPS treatment and during aging, consistent with previous findings (Clarke et al., 2018). On the other hand, we also observed notable differences in cell activation induced by aging and acute systemic inflammation, in both gene-expression and spatial patterns. In particular, the activation of microglia and astrocytes associated with the inflammation of oligodendrocytes in white matter of the corpus callosum were uniquely observed in the aging brain. Identifying the detailed molecular mechanisms underlying these commonalities and differences will require further mechanistic investigation of the roles of specific cytokines and other signaling pathways in the brain. Indeed, these two processes may intersect: intrinsic aging-related degenerative processes within the brain, such as degenerating myelin, which locally disrupt tissue homeostasis, may prime cells into a pro-inflammatory state, which could then be exacerbated by systemic factors (Pluvinage and Wyss-Coray, 2020).

The functional consequences of the disruptions to non-neuronal cellular homeostasis on neural circuits remain to be investigated. Many of the genes that we observed to be upregulated during aging, such as interleukins and complement proteins, have been shown to play a crucial role in regulating neural circuit organization and function via interactions between neurons and non-neuronal cells during development (Stevens et al., 2007; Vainchtein et al., 2018). Our cell atlas of the aging brain could facilitate future studies aiming to determine whether spatially localized upregulation of these molecules with age in turn causes localized disruptions to neural circuit function. Integrating these functional studies in mice with spatial transcriptomic measurements in humans in multiple conditions (normal aging, acute brain injuries, as well as chronic neurodegenerative disorders) will reveal how the inflammatory activation of non-neuronal cells contributes to cognitive impairment associated with advanced age and diseases at the neuronal and circuit levels.

## ACKNOWLEDGEMENTS

We thank members of the Zhuang and Dulac labs for helpful discussions. This work is in part supported by the Star-Friedman Challenge for Promising Scientific Research Award and the William F. Milton Fund Award. W.E.A. is a Junior Fellow of the Harvard Society of Fellows. C. D. and X.Z. are Howard Hughes Medical Institute Investigators.

## AUTHOR CONTRIBUTIONS

W.E.A, C.D., and X.Z. designed the experiments. W.E.A, T.R.B., and Z.A.S. performed experiments. W.E.A. analyzed the data. W.E.A, C.D., and X.Z. interpreted data and wrote the manuscript.

## STAR METHODS

- **KEY RESOURCE TABLE**
- **RESOURCE AVAILABILITY**

- Lead Contact
- Materials Availability
- Data and Code Availability
- **EXPERIMENTAL MODEL AND SUBJECT DETAILS**

- Animals
- **METHOD DETAILS**

- Single-nucleus RNA-sequencing
- snRNA-seq data analysis
- Gene selection for MERFISH measurements
- Design and construct of MERFISH encoding probes
- MERFISH encoding probe library and readout probe preparation
- Tissue sample preparation for MERFISH
- MERFISH imaging
- MERFISH data processing
- Integrated clustering analysis of the MERFISH and snRNA-seq data
- Imputation of gene expression
- LPS injection experiment
- **QUANTIFICATION AND STATISTICAL ANALYSIS**

- Brain region segmentation
- Gene module analysis
- Aging-related analysis
- Cell-cell proximity analysis
- Analysis of MERFISH data obtained from LPS-treated mice

### Key resources table

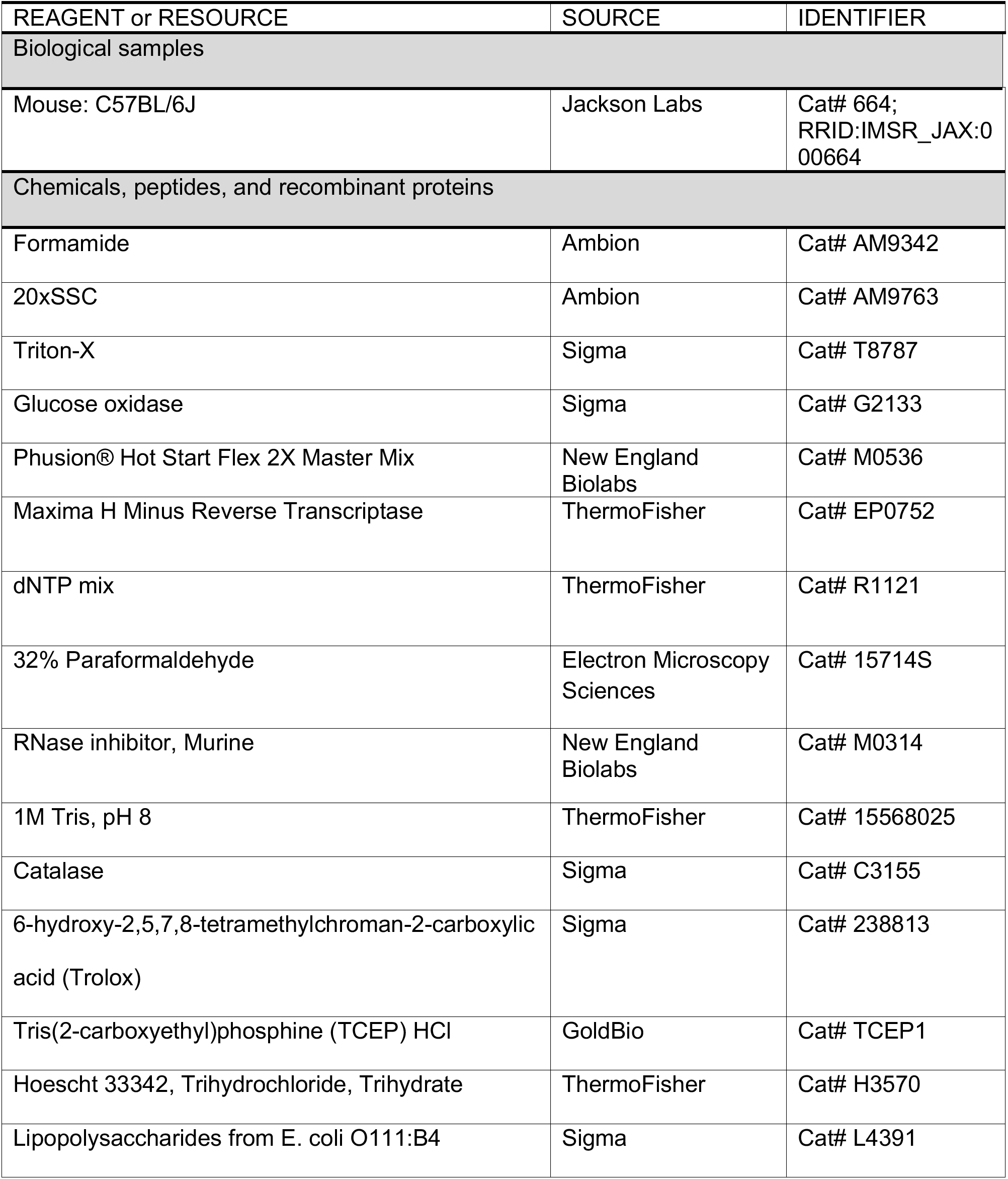
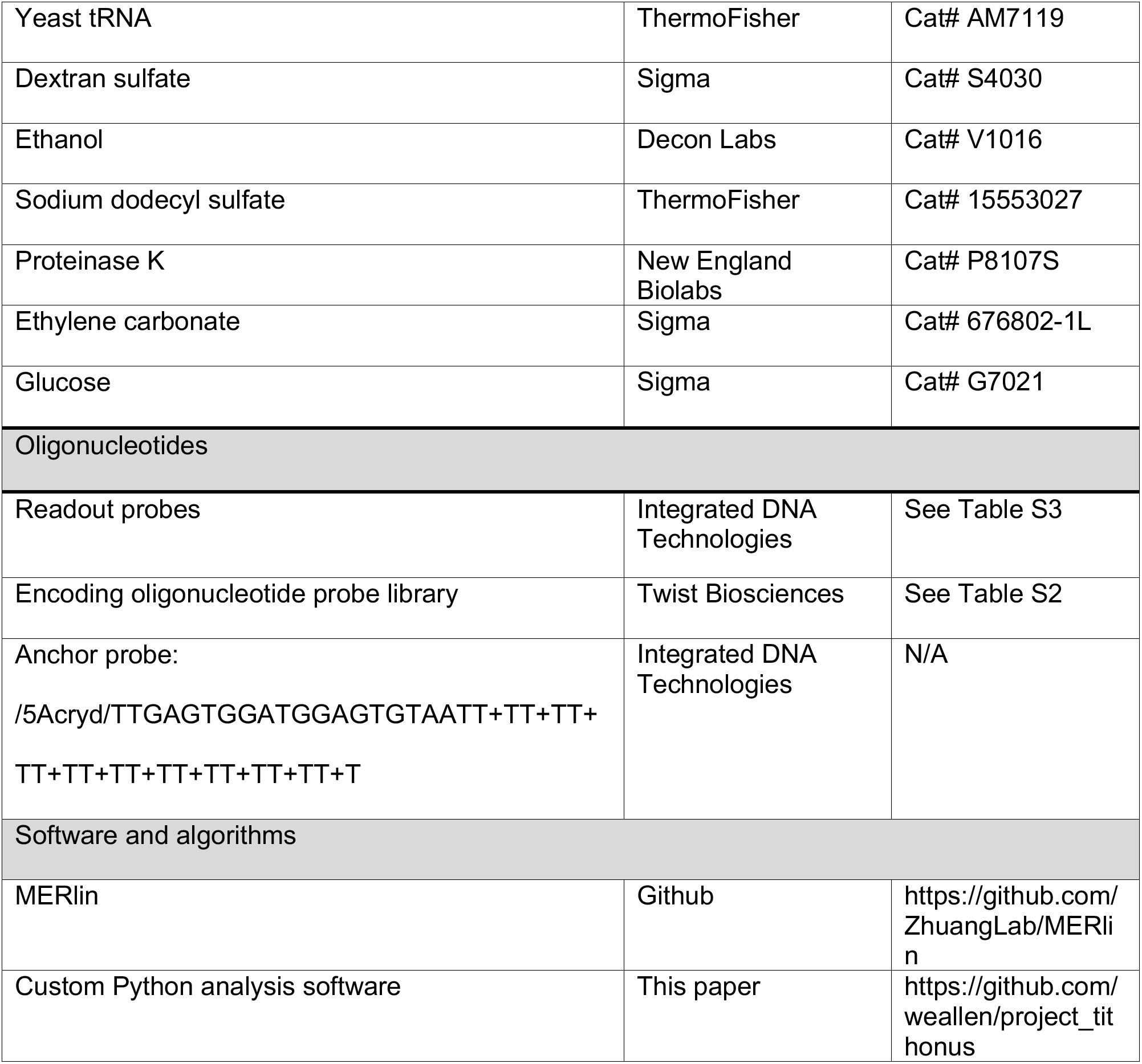

### RESOURCE AVAILABILITY

#### Lead Contact

Requests for resources and reagents should be directed to the lead contact, Xiaowei Zhuang.

#### Materials Availability

Oligonucleotide encoding probe sequences used for MERFISH imaging can be found in Table S2. Oligonucleotide readout probe sequences used for MERFISH imaging can be found in Table S3. These probes or templates for making these probes can be purchased from commercial sources, as described in the Key Resources Table.

#### Data and Code Availability

Single-cell RNA-seq data have been deposited to NCBI GEO data repository (GSE207848). All original code used in this work is available at: https://github.com/weallen/project_tithonus.

### EXPERIMENTAL MODELS AND SUBJECT DETAILS

#### Animals

Female C57BL6/J mice were used in this study. Mice were obtained from Jackson Laboratory at an age one week younger than the target age for sacrifice (4-week, 24-week, 90-week postnatal), and then housed at Harvard University Animal Facility for 1 week to acclimate before sacrifice. Mice were maintained on a 12 hr light/12 hr dark cycle (14:00 to 02:00 dark period) at a temperature of 22 ± 1°C, a humidity of 30–70%, with ad libitum access to food and water. Animal care and experiments were carried out in accordance with NIH guidelines and were approved by the Harvard University Institutional Animal Care and Use Committee (IACUC).

### METHOD DETAILS

#### Single-nucleus RNA-sequencing

Female mice aged 4 weeks or 90 weeks old were anesthetized with isofluorane and then acutely decapitated. Their brains were quickly harvested and cut into hemispheres and each hemisphere was frozen immediately on dry ice Optimal Cutting Temperature Compound (OCT, Fisher HealthCare) and then stored at –80°C until sectioning. Brains were taken from storage at –80°C and warmed to –18°C in a cryostat (Leica) for 20 minutes before sectioning. Sections were discarded until the beginning of frontal cortex was apparent. Brains were then blocked on the cryostat using a razor blade to a region containing frontal cortex and striatum. 100 µm coronal sections were then collected approximately from A/P +2 mm to A/P +1 mm, relative to bregma. The resulting sections were collected in an Eppendorft tube and stored at –80°C until snRNA-seq library preparation.

For snRNA-seq library preparation, nuclei were dounced in Nuclei EZ Prep nuclei extraction buffer (Sigma) + 1% RNase Inhibitor. Nuclei were then spun down, filtered through a 70 µm filter, stained with DAPI, and sorted on a FACS machine (BD FacsAria) to separate nuclei from debris. The resulting clean nuclei preparation were then counted and encapsulated on a 10X Genomics Chromium machine, using the 3’ Transcriptome V3.1 kit (10X Genomics). After encapsulation, the resulting libraries were reverse transcribed, amplified as cDNA, fragmented, and amplified as a final library following the manufacturer’s instructions. The resulting libraries were sequenced on a NovaSeq S4 flowcell (Illumina) to a target depth of ∼50,000 reads per nucleus.

#### snRNA-seq data analysis

Raw reads were mapped to the mm10 mouse reference genome and demultiplexed to generate a per-cell count matrix using CellRanger pipeline (10X Genomics). The resulting data were analyzed in Python using standard methods implemented in the package Scanpy. Briefly, putative doublets were first removed using Scrublet (Wolock et al., 2019). Cells with < 2,500 UMIs per cell and < 1,000 genes per cell were removed. Genes detected in < 3 cells were removed. Following standard procedures in Scanpy, per-cell counts were normalized to sum to 10^4^ counts per cells and log-transformed. A multi-level clustering approach was taken, where the cells were first clustered into major cell types then into clusters within those cell types as described in **Figure S1A**. Briefly, at each level highly variable genes were determined and included in the per-cell expression matrix, the total UMI number per cell and expression of mitochondrial genes were regressed out, and the resulting residuals were Z-scored. Principal component analysis was used to reduce the dimensionality to 50 principal components. A nearest neighbor graph was computed between cells using these principal components, and Leiden clustering was applied to separate the cells into distinct clusters.

First all cells were clustered into neurons and non-neuronal cells. Within the neurons, cells were clustered into inhibitory and excitatory neurons. Inhibitory neurons were further subclustered into medium spiny neurons (MSNs) and non-MSNs. Non-neuronal cells were subclustered into astrocytes, microglia, macrophages, oligodendrocytes, pericytes, vascular leptomeningeal cells (VLMCs). Each major cell type (excitatory, inhibitory, MSN, astrocytes, microglia, macrophages oligodendrocytes, pericytes, VLMCs) was then subclustered to obtain the final list of cell clusters.

#### Gene selection for MERFISH measurements

Genes were selected for MERFISH using a combination of automated and manual approaches. First, to identify age-related genes, linear regression was used to identify genes that were differentially expressed between two different ages (4-week and 90-week postnatal) in individual cell types or clusters determined by snRNA-seq. Briefly, using statsmodels, a Generalized Linear Model with a Negative Binomial link function was fit to the log-transformed UMI counts per cell for each gene *y_i_*:

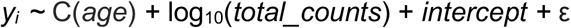

where C(*age*) is a binary categorical variable with the 4-week value set to be the reference level (i.e. C(4-week) = 0) and the C(90-week) value determined from the model. This model was fit separately for each cell type or cluster, which was compared with a null model that only accounts for technical variation in the total molecule counts per cell:

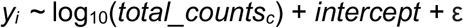

A likelihood ratio test was then computed between the full and reduced models to determine a *P*-value. These *P*-values were corrected for multiple hypothesis testing across all genes in all cell types or clusters to give the FDR-adjusted *P-*values, and genes with an FDR-adjusted *P*-value < 10^-6^ were considered. For each cell type and cluster, the genes differentially expressed between the two ages were sorted by the fitted C(90-week) value, and the top *N* genes with at least C(90-week) > 1.5 were included in the aging gene panel. For each major cell type, we included 5 genes and for each fine-leaflet cell cluster we included 2 genes. This approach attempts to balance the gene panel across all cell types and clusters, even if certain cell types or clusters may have more or fewer total numbers of differentially expressed genes with age.

To identify cell-type-marker genes, marker genes were identified for a particular cell population (cell type or cluster) using a one-vs-all approach. For each cell population, a *t*-test was performed for each gene between the cells within the cell population and all other cells not in that population. The resulting *P-*values were corrected for multiple hypothesis testing to give FDR-adjusted *P-*value. A gene was identified as a cell-type marker for a certain cell population if it satisfy the following conditions: i) it was expressed in at least 40% of cells within the specified cell population, ii) it had an FDR-adjusted *P-*value < 0.05, iii) it had a gene expression in the specified cell population that was at least two fold higher than the average expression in all cells not in that population, and iv) it was expressed in a fraction of cells within the specified cell population that was at least three times higher than the fraction of cells not in this population that expression the gene. Finally, the marker genes for each cell population were sorted by fold change in expression relative to the cell outside the cell population, and the top 15 marker genes for each cell population were then saved and used for marker selection. To select the final set of markers, we greedily added marker genes to the list such that each cell type or cluster had at least two marker genes included in the final gene panel.

In addition to these markers, known cortical layer markers (Zhang et al., 2021), genes related to microglial (Bohlen et al., 2019; Chen et al., 2020b; Keren-Shaul et al., 2017) and astrocyte (Clarke et al., 2018; Liddelow et al., 2017) activation, broad transcriptomic markers of aging (Benayoun et al., 2019), and markers for various immune cell types (T cells, B cells, macrophages) (Dong, 2021; Meizlish et al., 2021; Salvador et al., 2021) were curated from the literature and included. In total, 212 genes were included in the cell-type-marker gene panel and 204 genes were included in the aging gene panel.

#### Design and construction of MERFISH encoding probes

After the aging and cell-type-marker MERFISH gene panels were selected, a 20-bit code was created for the gene panels. Briefly, a 20-bit Hamming-weight 4 code was generated by first listing all possible combinations of 4 “on” bits embedded within 20 bits. This list was shuffled, and the first bit combination was randomly selected as the initial barcode. Additional barcodes were then added from this list by iterating through the other randomly shuffled barcodes and greedily adding to the codebook each additional barcode that had a Hamming Distance of at least 4 from all barcodes currently in the codebook. A collection of 500 such randomly sampled codes was generated, and each was scored on the total number of barcodes and the variance of number of barcodes utilizing each bit. A 20-bit, Hamming Distance 4 and Hamming-weight 4 code with the lowest variance that included at least 200 codewords was then used for both libraries, resulting in a 223-codeword final codebook.

Next, individual genes were assigned to barcodes in the codebooks. This assignment was initially random, then optimized to increase the average expected uniformity of the density of molecules per cell that were detected in each bit. This optimization was performed iteratively, using a simulated annealing algorithm to maximize the uniformity of expression across bits on average across all cell types.

First, we estimated the expected total number of molecules per cell for each bit as the sum of the expression (determined by snRNA-seq) of genes with barcode reading “1” at that bit. This was computed for each cell type, and the weighted average across cell types was computed weighted by the cell abundance in individual cell types. Individual gene assignments to barcodes were then swapped, and the average expression per cell per bit was recomputed. Assignment swaps that decreased the variance across bits were kept. This process was repeated until the algorithm converged when the variance stopped decreasing.

For each gene, we then designed a total of 92 encoding probes targeting that gene’s mRNA sequence. The encoding probes were designed as previously described (Moffitt et al., 2016a). Briefly, we selected regions with GC content between 30% and 70%, melting temperature T_m_ within 60-80 °C, isoform specificity index between 0.75 and 1, gene specificity index between 0.75 and 1, and no homology longer than 15 nt to rRNAs or tRNAs. For each library, each of the 20 bits was assigned to a 20-nt three-letter (A, T, C) readout sequence. Each encoding probe was constructed by concatenating the 30-nt target region with three 20-nt readout sequences for each probe. The readout sequences for each gene were randomly shuffled across all 92 encoding probes for that gene. The encoding probes additionally contained a 20-nt reverse transcription primer sequence at the 5’ end and a T7 promoter at the 3’ end, which also functioned as PCR primer sequences.

#### MERFISH encoding probe library and readout probe preparation

The encoding probe library was synthesized using large-scale arrayed oligo synthesis (Twist Biosciences) and then amplified, as previously described (Moffitt et al., 2016a). Briefly, the initial library was amplified using limited cycle PCR (Phusion Polymerase, NEB) monitored via qPCR. The library was then converted to RNA via *in vitro* T7 transcription (HiScribe T7 Quick High Yield Kit, New England Biolabs) from a T7 promoter integrated into the PCR product. The resulting RNA product was purified (RNA Clean and Concentrate, Zymo Research) and reverse transcribed (Maxima H– Reverse Transcriptase, ThermoFisher). The RNA in the resulting RNA:DNA hybrid was degraded using alkaline hydrolysis, and the final ssDNA product was first desalted via buffer exchange through a 7K MWCO desalting column (ThermoFisher) then concentrated using a phenol:chloroform extraction and ethanol precipitation, resulting in a 5-10 nM/probe final library.

For the 416-gene panel used in this study, 40 readout probes were used, each complementary to one of the 40 readout sequences on the encoding probes. For readout, the first twenty readout probes correspond to the 20 bits of the code used for the cell-type-marker gene panel and the remaining twenty readout probes correspond to the 20 bits of the code used for the aging-related gene panel. Each readout probe was conjugated to one of the two dye molecule (Alexa Fluor 750 or Cy5) via a disulfide linkage, as previously described (Moffitt et al., 2016a). The readout probes were obtained from Integrated DNA Technologies and resuspended immediately in Tris-EDTA (TE) buffer, pH 8 (Thermo Fisher), to a concentration of 100 µM, and stored at –20 °C until use.

#### Tissue sample preparation for MERFISH

Brains were prepared as described for snRNA-seq, with the addition of mice at the age of 24-week postnatal without LPS treatment, and mice at the ages of 4-week, 24-week, and 90-week postnatal following LPS injection. Sectioning was performed on a cryostat at –18°C. slices were removed and discarded until the frontal cortex and striatal target region was reached. In order to capture comparable sections across animals, starting from approximately A/P +2 mm relative to bregma, every other 10 µm section was captured onto a set of 6-8 lysine-coat silanized coverslips for MERFISH imaging, with each coverslip ultimately containing 3 to 4 individual sections. The coverslips were cleaned, silanized, and treated with poly-lysine as previously described (Zhang et al., 2021). The same anatomical region was identified for imaging post hoc after sample preparation, before the start of MERFISH imaging.

Tissue sections were fixed in 4% paraformaldehyde (Electron Microscopy Sciences) for 20 minutes, washed three times in 1× D-PBS, and then stored in 70% EtOH (Koptec) at 4°C for at least 18 hours to permeabilize the tissue. Tissue slices from the same mouse were cut at the same time and distributed onto 6-8 coverslips; multiple mice were sectioned at the same time. Coverslips were stored in 70% EtOH for less than two weeks until each biological replicate was successfully imaged once.

The tissue sections were stained with MERFISH encoding probes as previously described (Moffitt et al., 2016b). Briefly, the samples were removed from 70% EtOH and washed with 2× saline sodium citrate (SSC) three times. We then removed excess 2× SSC by blotting with a Kimwipe and inverted the coverslip onto a 50 µl droplet of encoding probe mixture in a Parafilm-coated Petri dish. The encoding probe mixture contained approximately 1 nM of each encoding probe, 1 µM of polyA-anchor probe (Integrated DNA Technologies) in 2× SSC with 30% v/v formamide, 0.1% wt/v yeast tRNA (ThermoFisher), 10% v/v dextran sulfate (Sigma), and 1% v/v murine RNase inhibitor (New England Biolabs). The polyA-anchor probe containing a mixture of DNA and LNA nucleotides (/5Acryd/TTGAGTGGATGGAGTGTAATT+TT+TT+TT+TT+TT+TT+TT+TT+TT+T, where T+ is locked nucleic acid, and /5Acryd/ is 5′ acrydite modification) was hybridized to the polyA sequence on the polyadenylated mRNAs, allowing these RNAs to be anchored to a polyacrylamide gel as described below. The sample was then incubated for 36-48 hours at 37°C.

After hybridization, the samples were washed in 2× SSC with 30% v/v formamide for 30 min at 47 °C for a total of two times to remove excess encoding probes and polyA-anchor probes. Tissue samples were cleared to remove lipids and proteins that contribute fluorescence background, as previously described (Moffitt et al., 2016b) Briefly, the samples were embedded in a thin 4% polyacrylamide gel and were then treated with a digestion buffer of 2% v/v sodium dodecyl sulfate (Thermo Fisher), 0.5% v/v Triton X-100 (Sigma) and 1% v/v proteinase K (New England Biolabs) in 2× SSC for 36 – 48 hours at 37 °C. After digestion, the coverslips were washed in 2× SSC for 30 min for a total of four washes and then stored at 4 °C in storage buffer of 2× SSC, 1% v/v murine RNase inhibitor (New England Biolabs) before imaging.

#### MERFISH imaging

We used a custom-built imaging setup in this study as previously described (Xia et al., 2019). All buffers and readout-probe mixtures were flowed onto the sample using a home-built, automatic fluidics system as previously described (Xia et al., 2019). Briefly, the samples were stained with 1 µg/mL Hoechst 33342 (ThermoFisher) and then loaded into a commercial flow chamber (Bioptechs) with a 0.75-mm-thick flow gasket. The first MERFISH round, containing both the first two readout probes labeled with Cy5 and AlexaFluor 750, as well as a probe complementary to the polyA anchor labeled with AlexaFluor 488, were then stained on the microscope. For each hybridization round, the fluorescent probes were hybridized in a buffer containing 2× SSC, 10% v/v ethylene carbonate (Sigma), and 0.1% Triton X-100, and were diluted to a final concentration of 3 nM. The samples were stained for 15 min, and then washed with readout probe buffer. Finally, imaging buffer was flowed into the chamber. The imaging buffer consisted of 2× SSC, 10% w/v glucose (Sigma), 60 mM Tris-HCl pH8.0, ∼0.5 mg/ml glucose oxidase (Sigma), 0.05 mg/ml catalase (Sigma) 50 µM trolox quinone (generated by UV irradiation of Trolox, 0.5 mg/ml 6-hydroxy-2,5,7,8-tetramethylchroman-2-carboxylic acid (Trolox, Sigma), and 0.2% v/v murine RNAase inhibitor (New England Biolabs).

After the readouts for the first round were hybridized, the sample was imaging with a low magnification objective (CFI Plan Apo Lambda ×10, Nikon) with 405-nm illumination to produce a low-resolution mosaic of the sections in the Hoeschst channel. We used this mosaic to generate a grid of tiled fields-of-view (FOV) covering the relevant areas of frontal cortex and striatum. We then switched to a high-magnification, high-numerical aperture objective (CFI Plan Apo Lambda ×60, Nikon), and imaged each FOV with a 7-plane z-stack with 1.5 µm spacing between the adjacent z-planes to cover the entire 10 µm thickness of the tissue section. For each FOV, we took images in the 750-nm, 650-nm, 560-nm, 488-nm and 405-nm channels: one image of the orange fiducial beads (560-nm) at the bottom z-plane, which was used as a fiducial marker to register the position of each FOV across multiple rounds of hybridization. For each z-plane, we took images of the readout probes (Alexa Fluor 750 and Cy5, 750-nm and 650-nm respectively), polyA probes (488-nm), and Hoeschst nuclear DNA stain (405-nm).

After the first round of imaging, the dyes were cleaved from the readout probes by flowing 2.5 mL of cleavage buffer (2× SSC and 50 mM of Tris (2-carboxyethyl) phosphine [GoldBio]) and incubating for 15 min, which cleaved the disulfide bond linking the dye to the readout oligonucleotide. The excess TCEP was removed by washing with 1.5 mL of 2× SSC.

For subsequent rounds of imaging, the same steps were carried out using readout-probe mixture containing 3 nM of the appropriate Alexa Fluor 750- and Cy5-labeled readout probes for each round.

The two gene panels (cell-type-marker panel and age-related gene panel) were imaged back-to-back on the same tissue sections, each gene panel was imaged in 10 rounds with two readout probes per round to readout the 20 bits. Each experiment took approximately 24-36 hours to image the relevant fields of view from 2 – 4 coronal slices.

#### MERFISH data processing

Imaging data were uploaded to the Harvard FAS Research Computing cluster and decoded using our previously published MERlin pipeline (Xia et al., 2019) with modifications on cell segmentation as described below. This pipeline provides the gene identity and spatial coordinates of each decoded molecules. For cell segmentation, we used the ‘cyto2’ model CellPose (Stringer et al., 2021), a deep learning based cell segmentation algorithm. This was applied only to Hoeschst-stained nuclei in order to avoid incorrect segmentation of neighboring cells. Decoding molecules were then assigned to the segmented nuclei to produce a cell-by-gene matrix that list the number of molecules decoded for each gene in each cell.

The cell-type-marker gene panel and aging-related gene panel were decoded separately. The segmented nuclei determined in these two decoding runs were then aligned by identifying the mutual nearest neighbor nuclei that were within 5 µm of each other. The very small number of nuclei that did not have a mutual nearest neighbor within this distance cutoff were removed from the dataset, as potentially incorrectly segmented cells, and the remaining nuclei were each assigned a vector of gene expression counts that included decoding results from both decoding runs. During data analysis, we observed that the readout of bit #20 of the aging-related gene panel was consistently dim in the majority of experiments, suggesting that genes detected in this bit may have a reduced detection efficiency. We thus excluded from all subsequent analysis the 40 genes that should be detected in this bit (i.e. genes whose barcodes read “1” at this bit), although the major conclusions in this paper were not altered if we included these genes in the subsequent analysis.

After decoding, cells from all MERFISH experiments were combined into a single dataset. Putative doublets were removed using Scrublet. Cells were then filtered to remove all cells with < 20 molecule counts per cell or with < 5 genes detected per cell. Cells that had a volume < 100 µm^3^ or a volume > 3 times the median volume across all cells were removed. Each cell’s gene expression values were normalized by dividing by the volume of that cell. The total normalized gene expression was computed for each cell, and cells with total normalized expression in the top and bottom 2% quantile were removed. Finally, these normalized values were scaled such that the sum of gene expression values per cell was equal to 250. The gene expression values were then log-transformed and Z-scored.

#### Integrated clustering analysis of the MERFISH and snRNA-seq data

The MERFISH expression matrix was concatenated with the normalized, log-transformed, and Z-score snRNA-seq expression matrix, which was subseted to include only the genes in the MERFISH gene panels. These data were then subjected to a two-step data integration and batch correction process, to first correct for modality-specific bias then for batch-specific bias using Harmony (Korsunsky et al., 2019) and BBKNN (Polański et al., 2020) respectively. First, Principal component analysis was performed on the join data matrix. Harmony was then used to adjust the principal components for modality-specific (MERFISH or snRNA-seq) effects, producing an integrated representation in the principal component space. Second, these integrated principal components were then used by BBKNN to compute a batch-corrected nearest neighbor graph, where each batch was an experimental run of MERFISH or snRNA-seq. This batch corrected nearest neighbor graph was subsequently used to further reduce the dimensionality of the dataset via UMAP or to compute integrated clusters via Leiden clustering.

To arrive at the final set of clusters, a semi-automated multi-level clustering approach was performed. Similar to the clustering approach used for snRNA-seq alone, cells were first clustered into neurons and non-neuronal cells. The neurons were then subclustered into excitatory and inhibitory neurons, and the inhibitory neurons were subclustered into MSNs and non-MSNs. The non-neuronal cells were clustered into oligodendrocytes, OPCs, astrocytes, microglia, vascular cells, and immune cells. These major cell types were then subclustered to arrive at the final list of clusters. Briefly, for each major cell type, the Harmony-corrected principal components were used via BBKNN to compute a batch-corrected nearest neighbor graph. This nearest neighbor graph was then used to perform Leiden clustering and to compute a UMAP plot for each major cell type. For each major cell type, differential gene expression was computed between each pair of subclusters using a *t*-test and the spatial locations of each cluster were plotted for manual inspection. The few clusters that appeared over-segmented based on heuristic criteria (no unique differentially expressed genes between the two clusters, largely overlap in UMAP space between the two clusters, a cluster with a very small cell number intermingled with a cluster with larger cell number in UMAP space) were then manually merged. The final set of clusters were annotated based on comparison of their key marker genes and/or spatial locations with previously annotated datasets (Chen et al., 2021; Tasic et al., 2018; Zeisel et al., 2018; Zhang et al., 2021).

#### Imputation of Gene Expression

To impute the genome-wide expression profiles of the cells measured by MERFISH, the gene expression profiles of the snRNA-seq cells most similar to each cell measured with MERFISH were averaged together. This computation used the PCA embedding produced through Harmony integration to identify the nearest neighboring snRNA-seq cells for each MERFISH cell, using the top 30 principal components in the jointly embedded PCA decomposition. The genome-wide expression profiles of the 10 nearest neighboring snRNA-seq cells for each MERFISH cell was averaged together to produce the imputed gene expression profile for that MERFISH cell.

#### LPS Injection Experiment

Female C57BL6/J mice were injected intraperitoneally with 0.5 mg/kg lipopolysaccharide (LPS) derived from *Escherichia coli* O111:B4 (Sigma) diluted in PBS at Zeitgeiber Time 9. Animals were euthanized 24 hours after injection and brains harvested for MERFISH analysis. LPS was titrated following reconstitution to optimize dosage and ensure consistent potency across experiments.

### QUANTIFICATION AND STATISTICAL ANALYSIS

#### Brain region segmentation

In order to automatically segment anatomical regions across the many individual sections included in the MERFISH experiment, we developed a semi-supervised method to cluster cells based on the cell type composition of their local spatial neighborhood. For each cell, we computed the abundance of cells from all clusters within a 100 µm radius of this cell, presented in the form of a vector with N_cluster_ dimensions, where N_cluster_ in the number of clusters. We then combined these vectors across all cells to form an N_cell_ x N_cluster_ matrix, where N_cell_ is the number of cells. We applied principal components analysis to this matrix to reduce the dimensionality of this matrix, and then applied *k*-means clustering to segment the cells into *k* = 20 spatial clusters. This produced an overclustered segmentation of cells in space. We then hierarchically ordered these spatial clusters and manually merged spatial clusters that were near each other in the cluster hierarchy and appeared over-segmented when their spatial profiles were plotted, to arrive at a final set of 8 spatial clusters, which map closely to known anatomical structures, including the pia, cortical layer 2/3 (L2/3), cortical layer 5 (L5), cortical layer 6 (L6), corpus callosum, striatum, ventricle, and subcortical olfactory areas.

#### Gene Module Analysis

Gene modules were identified from scRNA-seq data by first grouping cells from each major cell type. For each group, variable genes were then selected and a gene-gene correlation matrix was computed by taking the first 50 singular values from the singular value decomposition of the gene expression matrix and computing the dot product of it with its transpose. This gene-gene correlation matrix was then Z-scored and Z-scores less than 0.1 were set to zero to sparsify the matrix. Each gene in this matrix was then reduced to 2 dimensions using UMAP. Genes were then clustered into modules in this reduced dimensional space using the DBSCAN clustering algorithm. To remove modules that were not associated with any genes in a statistically significant manner, we compared the mean gene-gene correlation per module under this clustering with a shuffled distribution where the module identities of the genes were randomly permuted 1000 times to determine the *P*-value. These *P*-values were then FDR corrected and any modules with FDR < 0.1 were removed.

#### Aging-related analysis

Activation scores were computed from the normalized, log-transformed, Z-scored gene expression values using the score_genes function in Scanpy, which computes the summed expression value of a set of genes minus the average expression of randomly selected background genes. The genes used for astrocyte activation were: *C4b, C3, Serpina3n, Cxcl10, Gfap, Vim* (Clarke et al., 2018; Liddelow et al., 2017; Zamanian et al., 2012). The genes used for microglial activation were: *B2m, Trem2, Ccl2, Apoe, Axl, Itgax, Cd9, C1qa, C1qc, Lyz2, Ctss* (Bohlen et al., 2019; Chen et al., 2020b; Keren-Shaul et al., 2017). The genes used for oligodendrocyte activation were: *C4b*, *Il33, Il18,* which were identified based on differential gene expression of oligodendrocytes over aging.

To compute the activation scores of microglia or astrocytes as a function of distance to another cell type, the average activation scores of all microglia or astrocytes were computed as a function of distance from a reference cell type (Oligo, Endo, or VLMC). For each individual astrocyte or microglial cell, the nearest neighbor of a particular comparison cell type was identified within a radius of 80 µm of that astrocyte or microglia using a *k*D-tree search implemented in scikit-learn. The distance from the astrocyte or microglia to that comparison cell type was saved along with that astrocyte or microglia’s activation score. The average activation score for all microglia or astrocytes were computed at 1 µm stepping from 0 to 80 µm, with a sliding 30-µm-wide window. Finally, the mean activation across this range was subtracted.

#### Cell-cell proximity analysis

To compute the proximity frequency between two sets of cells, i.e. cell type *A* and cell type *B*, first the true cell-cell proximity frequency µ_true_ was computed as follows: for each cell in cell type *A,* the average number of cells of cell type *B* were counted within a radius of 30 µm. To compute the null distribution of proximity (the probability that two cell types would be within 30 µm just by chance, given their local density), the locations of each cell of cell type *B* were then randomly jittered independently in both spatial dimensions (x and y) using a uniform distribution over the interval (–100 µm, 100 µm), and the number of cells in cell type *B* within 30 µm of each cell in cell type *A* was re-computed. This was repeated 1,000 times, to form a background distribution of the frequency of cell-cell proximity that would be expected to occur due to chance. The enrichment of cell-cell proximity between cell type *A* and cell type *B* was then computed as the log of the ratio between the true proximity frequency µ_true_ and the average background frequency µ_background_: log_2_(µ_true_/ µ_background_). A *P-*value for this fold change was found by computing the Z-score distribution across the 1,000 randomizations:

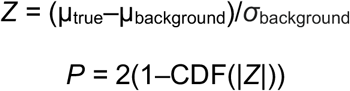

The enrichment and *P-*value were computed for each pair of major cell types, and the resulting *P-*values were FDR adjusted across all cell type pairs.

#### Analysis of MERFISH data obtained from LPS-treated mice

To transfer cell type and cluster labels obtained from cells in the non-LPS-treated mice to cells from mice treated with LPS, we took a supervised classification approach. First, the +LPS data were pre-processed as described earlier to remove doublets and obtain normalize, log-transformed, and Z-scored expression values. The –LPS and +LPS MERFISH data were then co-embedded in a joint principal component space using Harmony to compute the first 25 principal components. A multilayer perceptron classifier from scikit-learn was trained on the cell type and cluster annotations from the –LPS cells, using the first joint principal components as input. The classifier was then applied to the +LPS cells, to yield a final set of predicted cluster annotations that were used for subsequent analysis.

To identify genes differentially expression between the +LPS and –LPS conditions, for each gene in the MERFISH library, a model was fit using ordinary least squares that compared the two conditions for young (4-week) mice. For each gene *i*, the average expression was modeled for cells in a given cell type in the +LPS and –LPS conditions using the following ordinary least squares model, fit to the normalize, log-transformed, Z-scored MERFISH expression data:

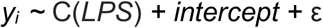

where C(*LPS*) is a categorical variable with the –LPS value set to be the reference level (i.e. C(– LPS) = 0) and the C(+LPS) value determined from the model. This was then compared with a null model lacking the C(*LPS*) categorical variable, i.e. *y_i_ ∼ intercept +* ε, to determine the gene’s FDR-adjusted *P-*value. Likewise, we performed similar analysis to determine genes differentially expression between the 4-week-old and 90-week-old mice without LPS treatment from the MERFISH data. To identify whether a gene was upregulated by LPS-only, aging-only, or by both LPS and aging, the C(+LPS) or C(90-week) values for each gene for each cell type were computed. These values were then Z-scored across genes, and genes with a Z-score > 2 and FDR-adjusted *P-*value < 0.05 in a particular condition was determined as being significant for that condition.

## Supplementary Figures

**Figure S1:**
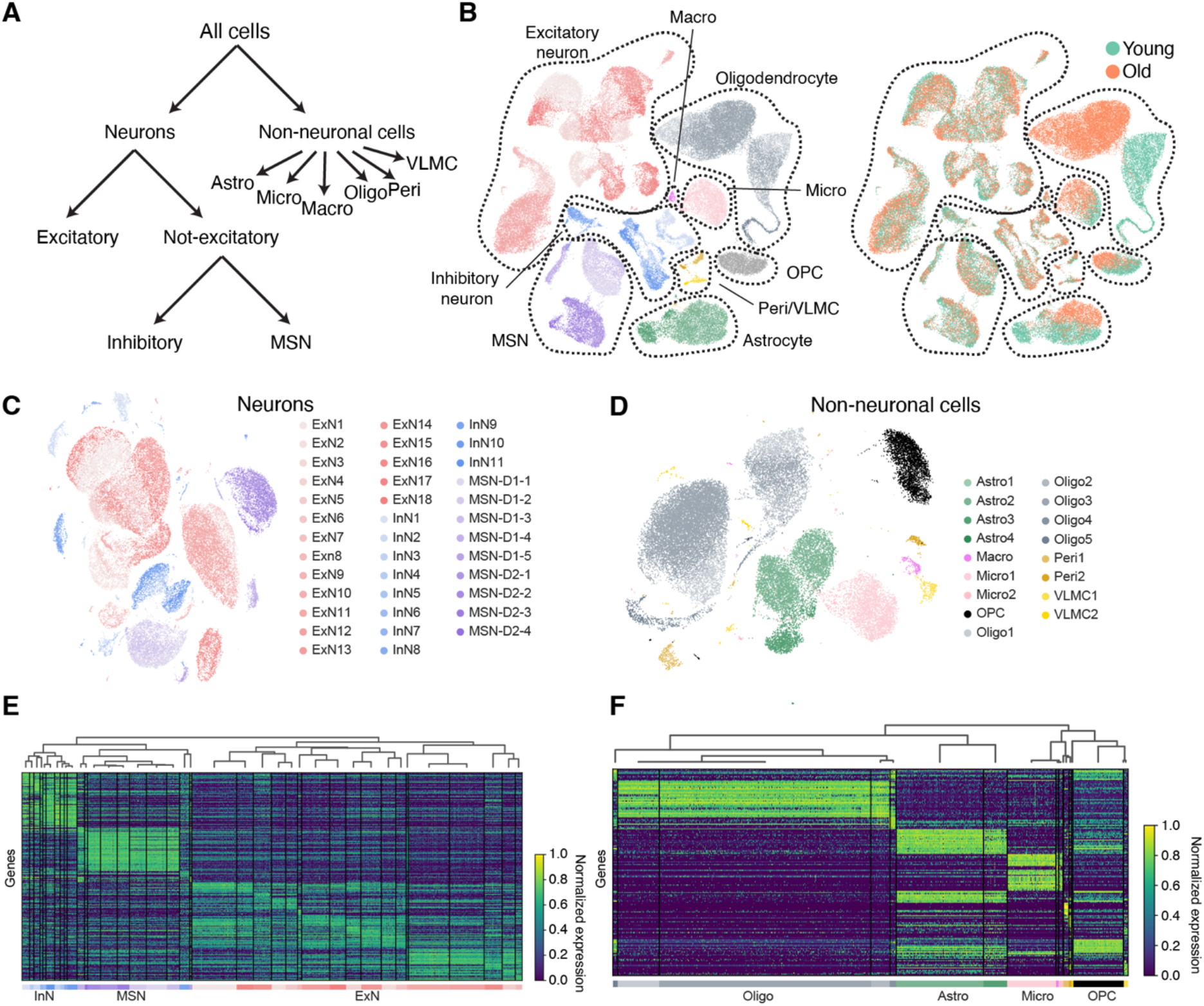
Clustering of cell types measured by snRNA-seq. **(A)** Flow chart for multi-step cell clustering analysis. First, cells were divided into neurons and non-neuronal cells. Neurons were then divided into excitatory and non-excitatory neurons, and the latter were then divided into medium spiny neurons (MSN) and other inhibitory neurons. Each of these major types were further divided into finer clusters. Non-neuronal cells were divided into astrocytes (Astro), microglia (Micro), macrophages (Macro), oligodendrocytes (Oligo), pericytes (Peri), and vascular leptomeningeal cells (VLMC), which were further divided into finer clusters. **(B)** UMAP visualization of cells colored by cell types (left) or ages (right). (**C, D**) UMAP plots of neuronal clusters (C) and non-neuronal clusters (D). (**E, F**) Heatmap of top differentially-expressed marker genes across different neuronal clusters (E) and non-neuronal clusters (F). Marker genes were determined using a *t*-test, with FDR-adjusted *P*-value < 0.05 and a fold change > 1.5. Expression values are normalized to the maximum value within each row.

**Figure S2:**
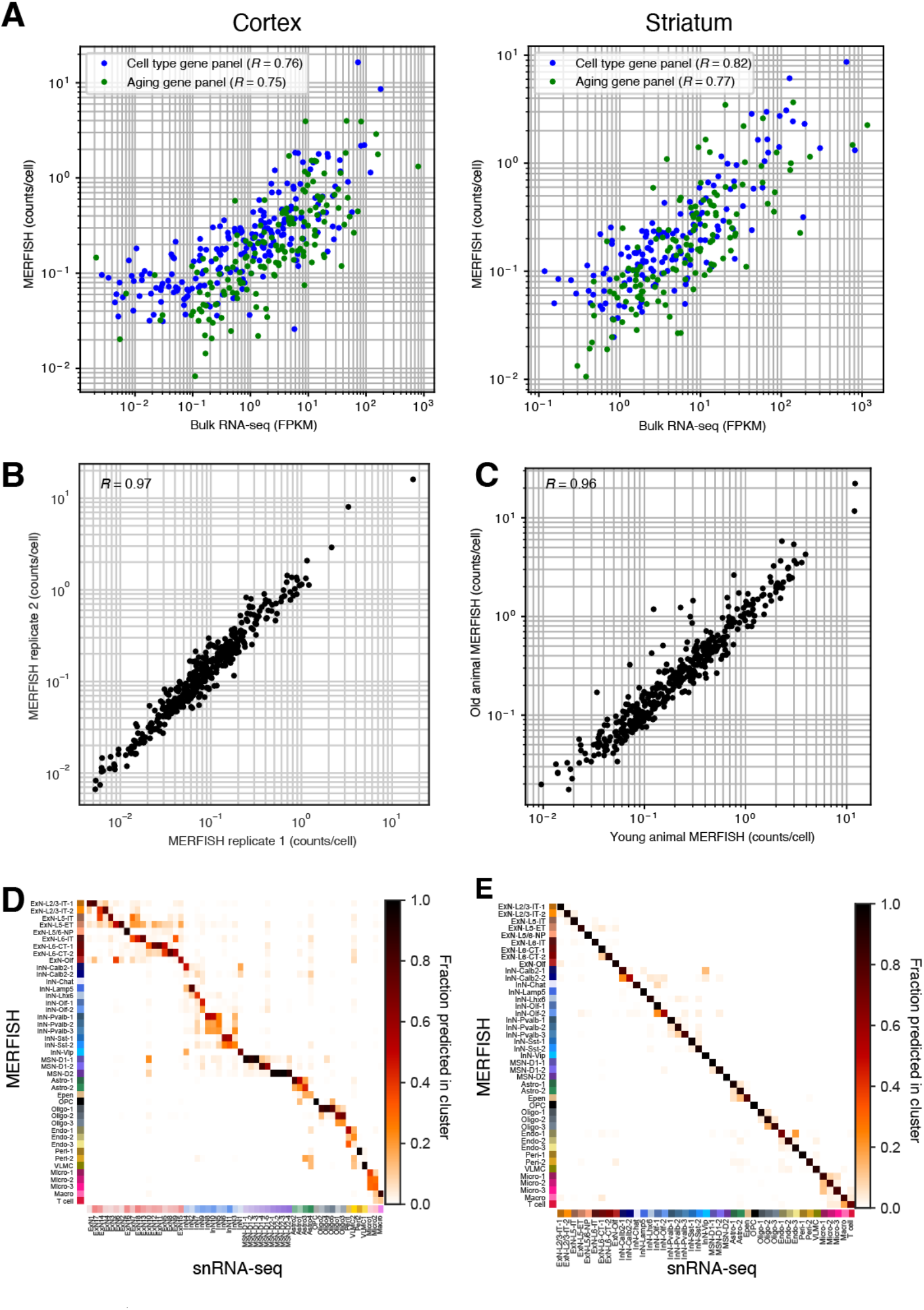
Correlation of MERFISH measurement with bulk RNA-seq, between MERFISH replicates, and with snRNA-seq. **(A)** Correlation of average expression values of individual genes included in the cell-type marker gene panel and aging-related gene panel measured by MERFISH with FPKM values of the same genes measured by bulk RNA-seq for motor cortex (left) and striatum (right). Pearson correlation coefficients *R* are given in the plots. **(B)** Correlation of average expression values of individual genes between two biological replicates measured by MERFISH. Pearson correlation coefficient *R* is given in the plot. **(C)** Correlation of average expression values of individual genes between all replicates from young mice and all replicates from old mice measured by MERFISH. Pearson correlation coefficient *R* is given in the plot. **(D)** Correspondence between MERFISH clusters, determined from an integrated snRNA-seq and MERFISH clustering analysis shown in **Figure 1**, and snRNA-seq clusters, determined from a separate clustering analysis of the snRNA-seq data alone shown in **Figure S1**. **(E)** Correspondence between MERFISH and snRNA-seq clusters, both determined from the integrated MERFISH and snRNA-seq clustering analysis. In (D) and (E), a multilayer perceptron neural network classifier was used to predict a MERFISH cluster identity for each cell measured in the snRNA-seq dataset. The fraction of cells from any given snRNA-seq cluster (columns) that was predicted to have each MERFISH cluster label (rows) was plotted.

**Figure S3:**
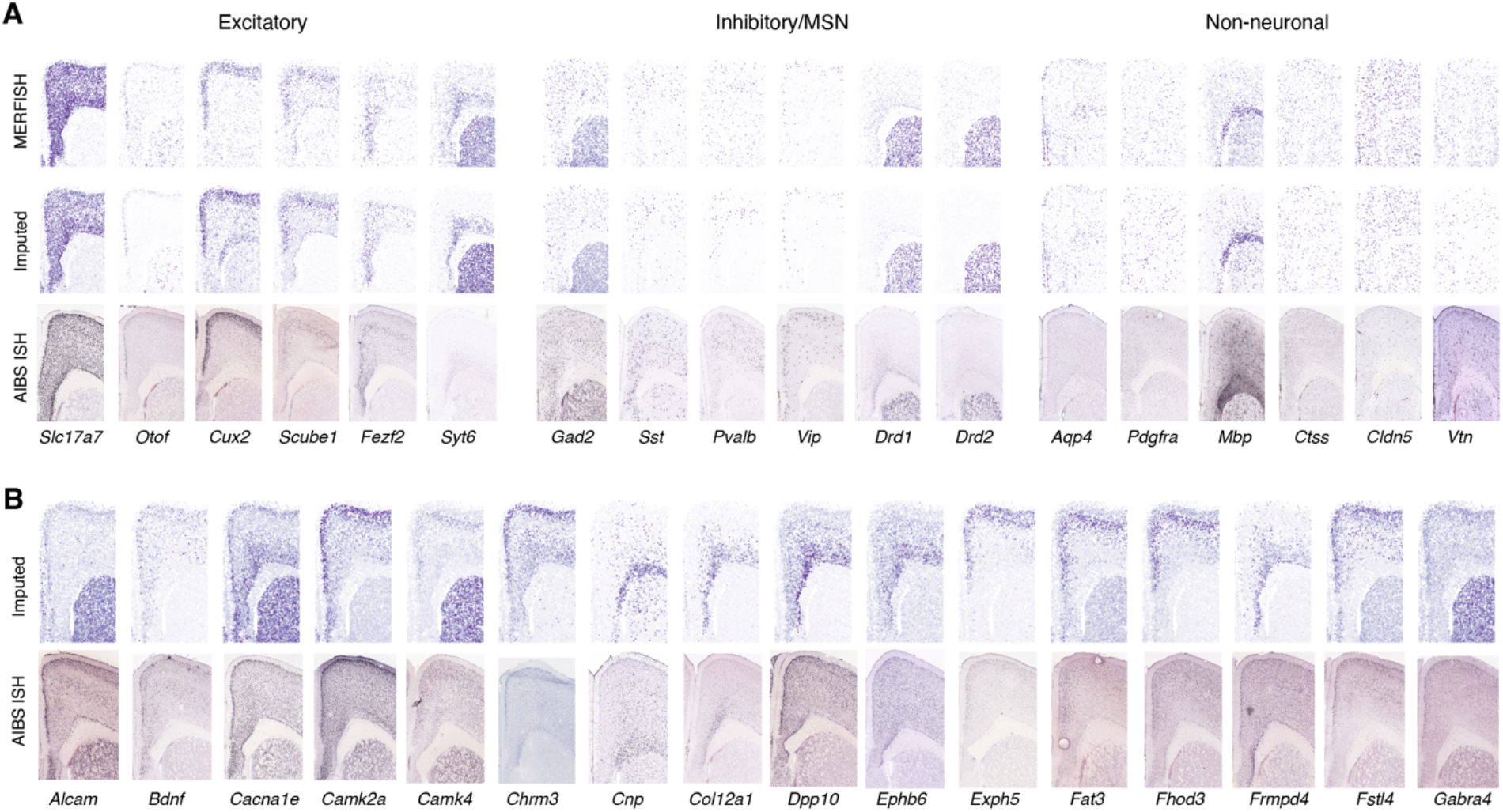
Comparison of imputed genes with MERFISH-measured genes and with Allen Brain Institute *in situ* hybridization data. **(A)** Spatial plot of expression of marker genes for different excitatory, inhibitory, and medium spiny neurons, and non-neuronal cell types in measured MERFISH data, imputed gene expression, and corresponding Allen Institute for Brain Science (AIBS) *in situ* hybridization (ISH) data. **(B)** Spatial plots of imputed genes that were not measured in the MERFISH gene panel and corresponding AIBS ISH data. Genes in (B) were randomly selected from imputed genes with clear spatial patterns. The AIBS ISH data in (A) and (B) are taken from https://mouse.brain-map.org/ (credit: Allen Institute),

**Figure S4:**
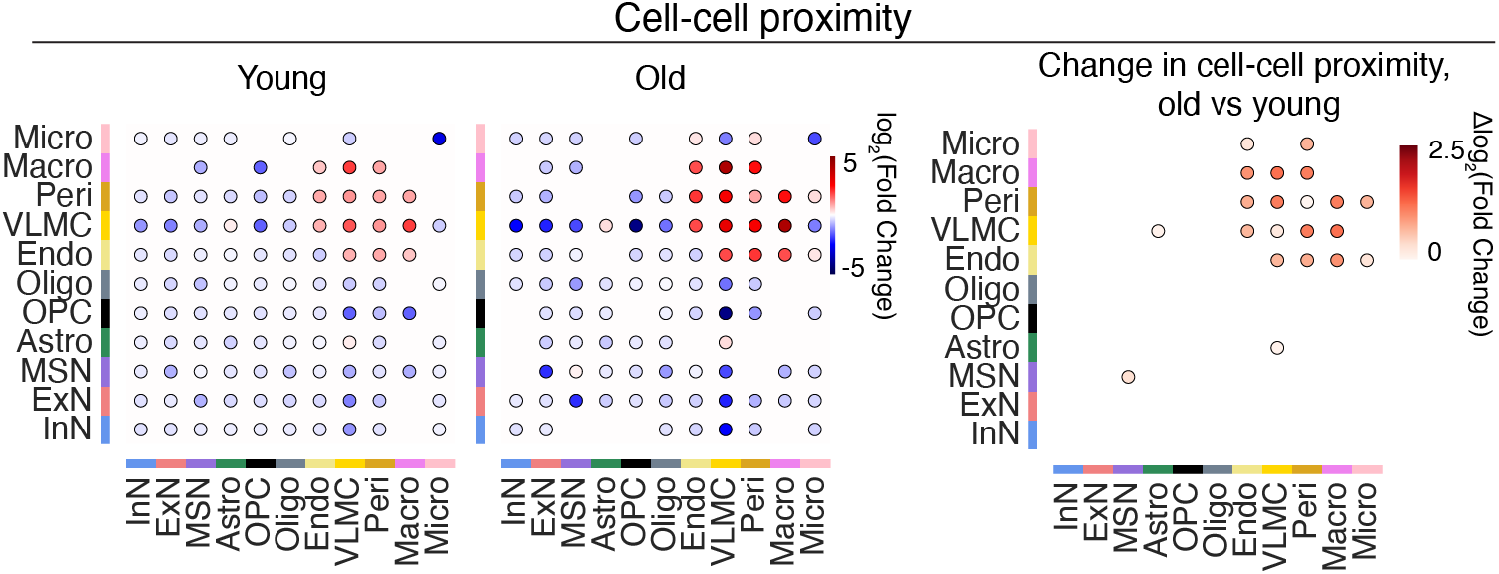
Cell-cell proximity between different cell types. (Left) Enrichment of cell-cell proximity between different cell-types. Enrichment in proximity between any give pair of cell types, A and B, was computed as the Log_2_ (Fold change) in the frequency of A-B cell pairs within a 30 µm radius, relative to the average A-B cell-pair frequency in a background distribution where each cell was randomly shifted by up to 100 µm to disrupt the original spatial relationship between neighboring cells. Plot shows only cell-type pairs with an FDR-adjusted *P-*value < 0.05. (Right) Difference in enrichment of cell-cell proximity between young and old animals.

**Figure S5:**
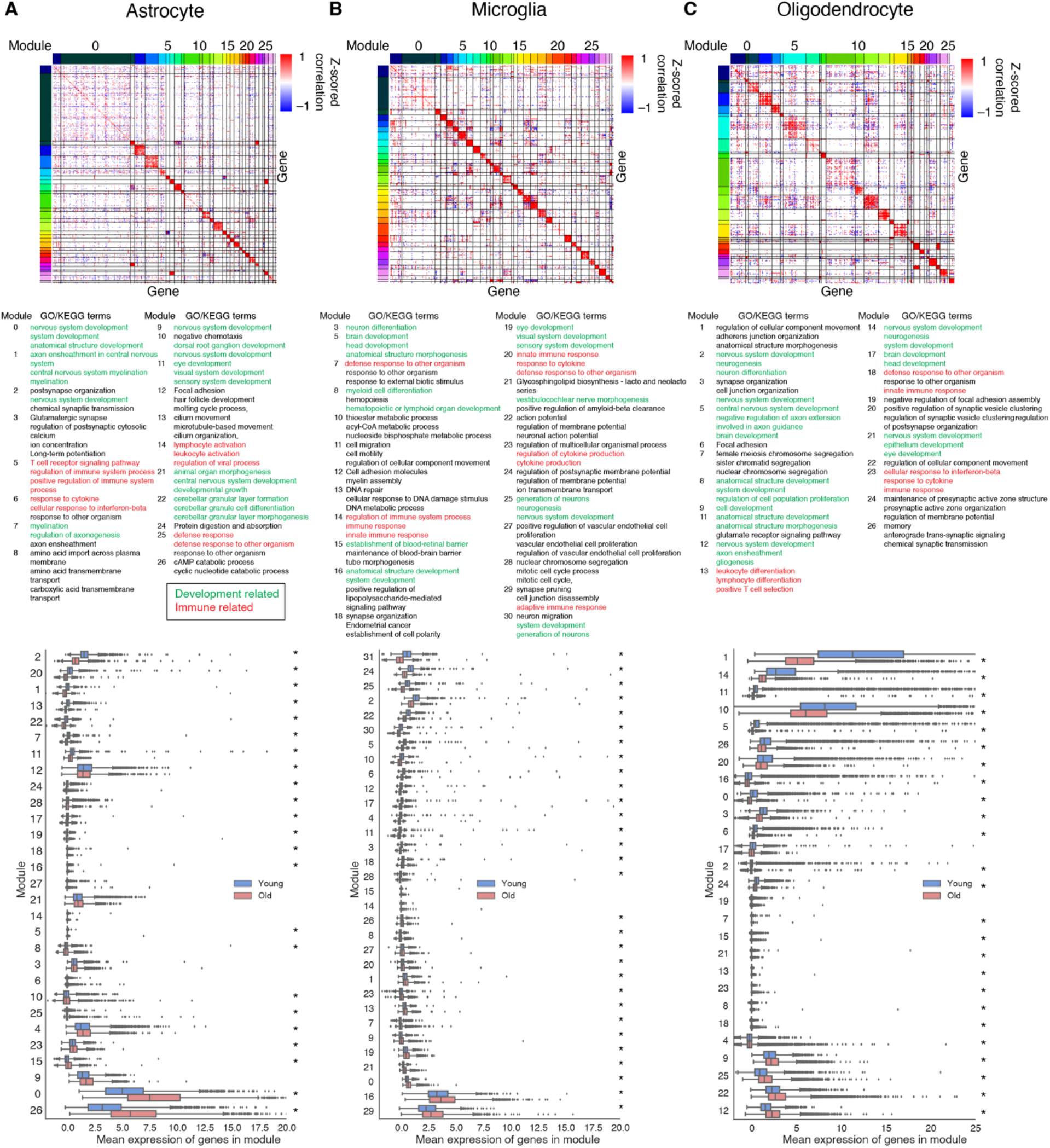
Gene modules analysis for astrocytes, microglia, and oligodendrocytes. (**A, B, C**) Top: Gene-gene correlation matrix for variable genes in astrocytes (A), microglia (B), and oligodendrocytes (C). Z-scored Pearson correlation coefficient for the expression levels of pairs of genes are shown and ordered by hierarchical clustering to show groups of genes with correlated expression, which we termed gene modules. Middle: Top Gene Ontology (GO) or Kyoto Encyclopedia of Genes and Genomes (KEGG) terms enriched in modules (up to three terms from the top five terms are shown per module). Only modules with any enriched terms are shown. Development-related terms are colored green and immune-related terms are colored red. Bottom: Expression levels for each module (calculated as the mean expression of the genes included in the module) in young and old mice, sorted by average difference between young and old. Box plots show the distribution of each module’s expression across individual cells within a cell type grouped by age. Box shows median and interquartile range, whiskers show minimum and maximum, outliers are shown as fliers. * FDR-adjusted *P*-value < 0.05, from independent *t*-test of module expression levels across cells between young and old mice.

**Figure S6:**
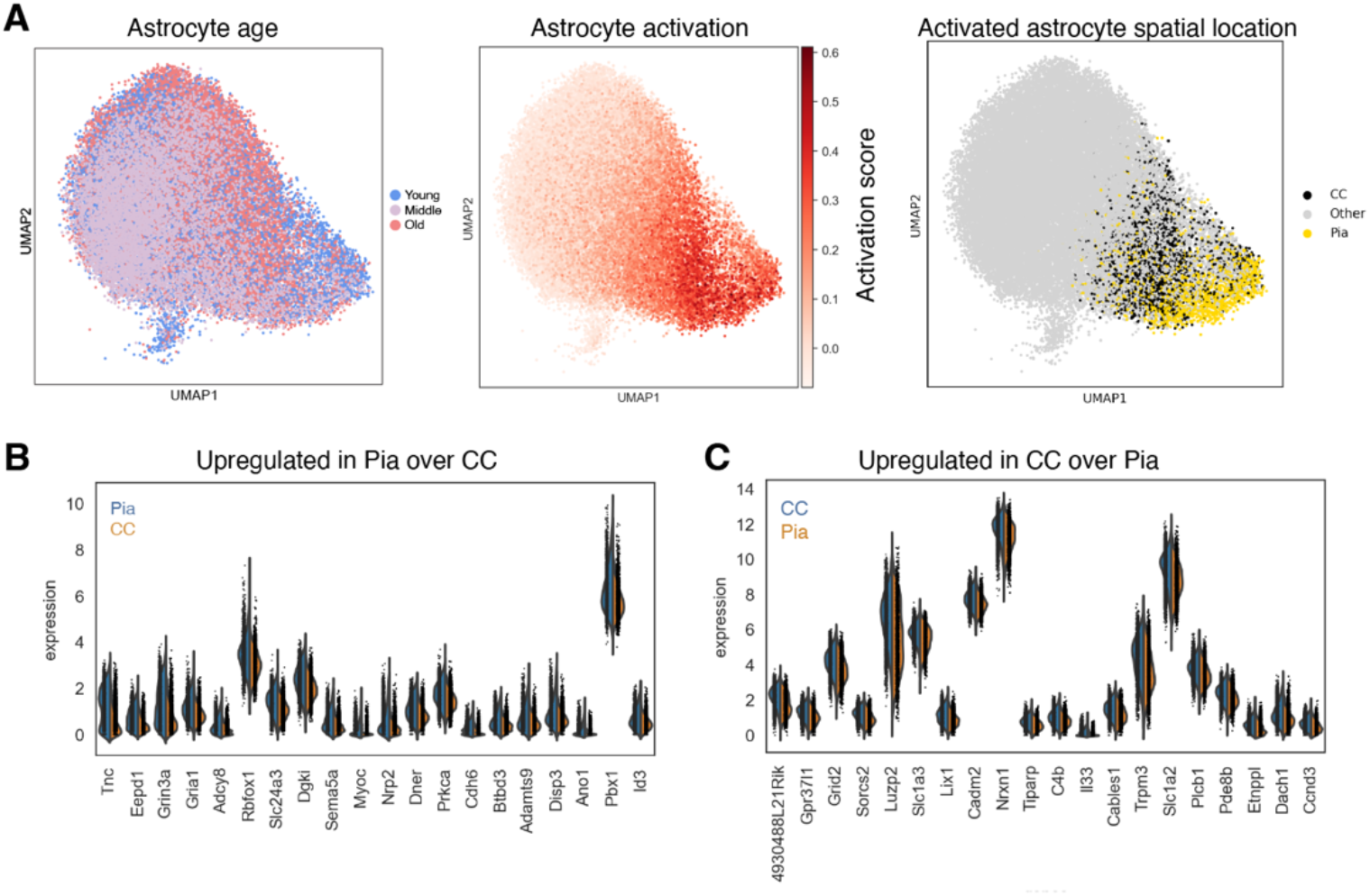
Molecular heterogeneity of activated astrocytes in pia and corpus callosum. **(A)** Left: UMAP plot of astrocytes colored by the age of cells. Middle: UMAP plot of astrocytes colored by the activation score of cells. Right: UMAP plot of astrocytes colored by the spatial location of activated cells (black: activated astrocytes in corpus callosum [CC]; Yellow: activated astrocytes in pia: Grey: all other astrocytes). (**B, C**) Relative expression levels of differentially expressed genes between activated astrocytes in pia vs corpus callosum, showing genes that are upregulated in pia relative to corpus callosum (B) and upregulated in corpus callosum vs pia (C). Activated cells were defined as having activation score greater than 1 standard deviation above the mean.

**Figure S7:**
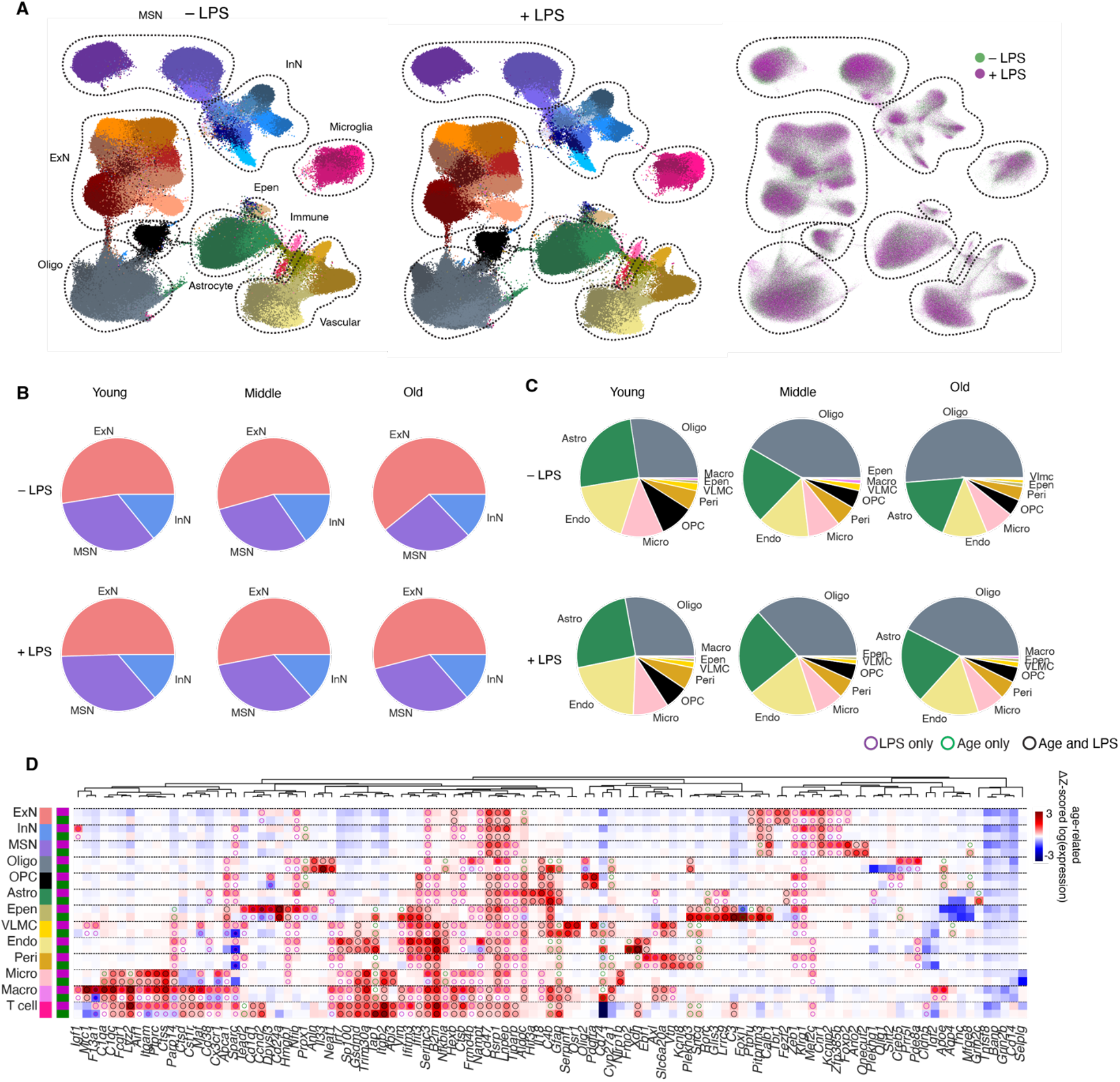
Comparison of cell-type compositions between –LPS and +LPS treatment conditions, and changes in gene expression during aging and in response to LPS treatment. **(A)** (Left) Visualization of clusters in an integrated UMAP space for cells measured in the –LPS condition and cells measured in the +LPS condition. Cells are colored by their cluster identities. (Right) Overlay of cells colored by –LPS or +LPS condition. **(B)** Pie chart of major neuronal cell-type composition across the three different ages, for –LPS and +LPS conditions. **(C)** Pie chart of major non-neuronal cell-type composition across three different ages, for –LPS and +LPS conditions. **(D)** Quantification of changes in expression of individual genes for each cell type designated on the left, where alternating rows show the change in Z-scored log(gene expression) for LPS-related changes (comparing +LPS vs. –LPS, young mice) and aging-related changes (comparing young vs old mice, –LPS). Black circles mark genes that are upregulated in both conditions, magenta circles mark genes upregulated in response to LPS treatment only, and green circles mark genes upregulated in aging only. Only genes with change in Z-scored log(gene expression) > 2 and FDR-adjusted *P*-value < 0.05 in at least one condition for at least on cell type are shown.

**Figure S8:**
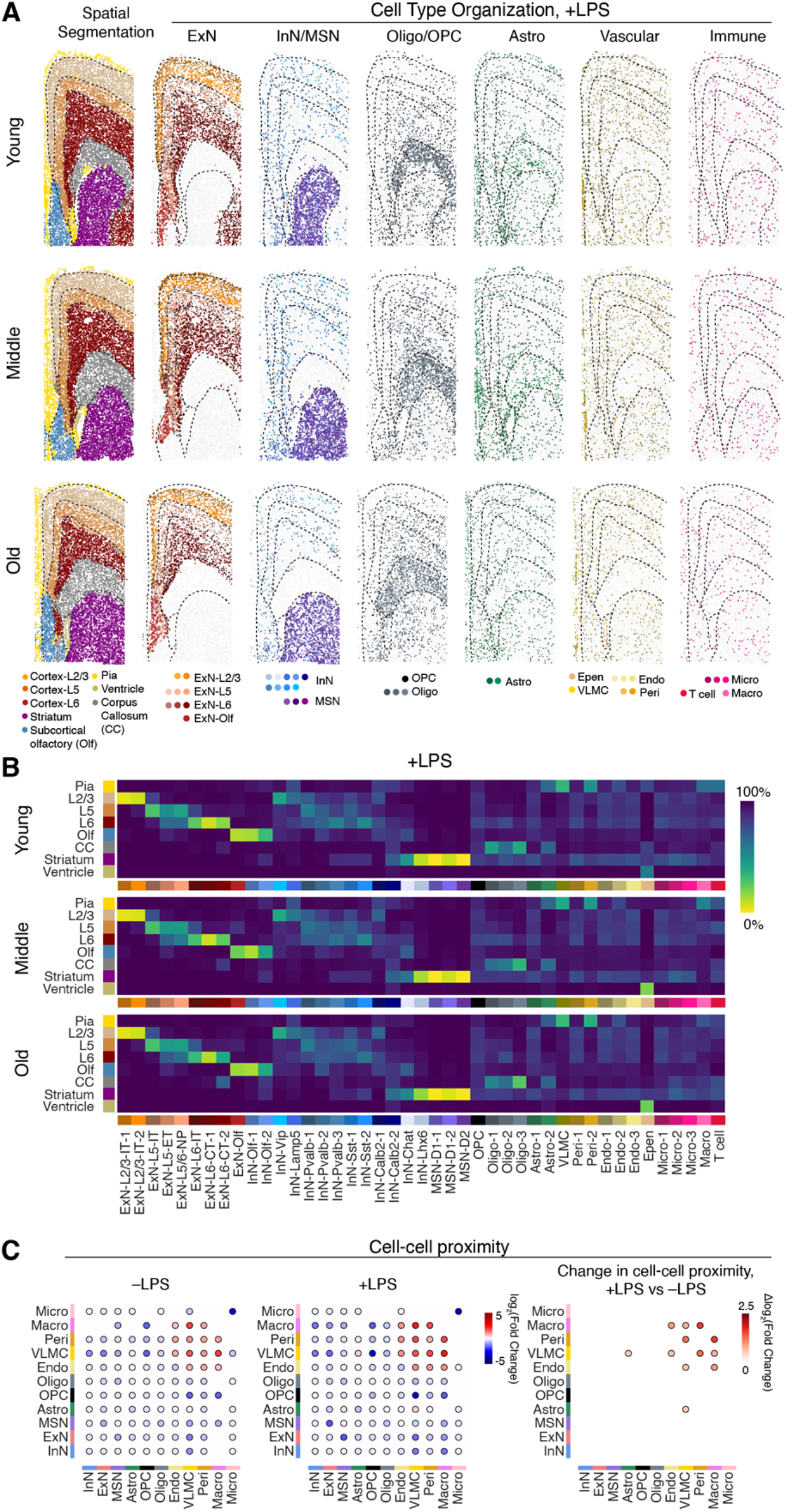
Spatial organization of cells after LPS treatment. **(A)** Visualization of spatial organization of different neuronal and non-neuronal cell types across three different ages in +LPS condition, as in **Figure 3A**. **(B)** Quantification of fraction of cells in different anatomical regions in +LPS condition across three different ages, as in **Figure 3B**. **(C)** (Left) Enrichment of cell-cell proximity between different cell types for –LPS condition and +LPS condition in young animals. Enrichment is defined as in **Figure S4**. (Right) Difference in enrichment of cell-cell proximity between –LPS and +LPS condition in young animals.

## Supplementary Table Captions

**Table S1:** MERFISH codebooks for the cell-type marker gene and aging-related gene panels. The “Celltype Codebook” sheet contains the codebook for genes that are cell-type markers and the “Aging Codebook” sheet contains the codebook for aging-related genes. The first column lists the gene names. The following columns list the binary values for each of the 20 bits and each bit is indicated by name of the corresponding readout sequence. Barcodes used as blank controls are denoted by a gene name that begins with “Blank-”.

**Table S2:** MERFISH encoding probes for the cell-type marker gene and aging-related gene panels. The “Celltype Encoding Probes” sheet contains encoding probes for genes related to cell type identity and the “Aging Encoding Probes” sheet contains encoding probes for aging-related genes. The targeted gene name and encoding probe sequence are provided for each encoding probe.

**Table S3:** MERFISH readout probes. For each readout probe, the bit number, readout probe sequence name, and readout probe sequence are provided.

